# Retinal resuscitation in post-mortem eyes

**DOI:** 10.64898/2026.06.25.733416

**Authors:** Eimear M. Byrne, Umberto Di Vicino, Nairouz Farah, Marta Cadevall Angles, Manuel Fernández Merino, Mikhail Rotkevich, Daniel Caravaca Rodriguez, Mark Balashov, Miguel Guerra Solano, Mohammed Salim Ibrahim, Jose Alejandro Marin Figuera, Javier José Puig Galy, Alberto Sandiumenge, Ricardo Casaroli Marano, Jack Lee, Yossi Mandel, Maria Pia Cosma

## Abstract

Vision loss compromises the quality of life of millions of people worldwide. Currently, vision-restoring therapies are lacking. Post-mortem preservation of human eyes may enable whole eye transplantation (WET) or therapeutic development. We developed an approach to cannulate the ophthalmic artery and perfuse post-mortem pig and human eyes using a custom-built device, Eye-in-Care-Box (ECaBox). Retinal vasculature imaging and reconstruction were used to assess perfusion efficacy using a supervised deep learning model. Without intervention, the retina degrades post-mortem, whereas perfusion preserves retinal structure and cell viability for up to 24-hours. Eyes were perfused within 30 minutes post-extraction, and regained light-responses persisted for up to 10 hours after death, challenging the notion that response to light ends at death. Our findings show that intact eyes can be resuscitated and preserved outside the body, supporting the ECaBox platform for WET preservation and pre-clinical therapeutic testing. Ultimately, ECaBox may bring vision-restoring treatments to patients.

## Introduction

The retina is a layered neural tissue in the posterior chamber of the eye, which converts light into neural signals that are transmitted to the brain. It has a high metabolic demand and avascular zones, and its function relies heavily on maintenance of structural integrity^1^. Thus, maintaining the retina inside the intact eye outside the body involves unique vascular and metabolic challenges compared to other organs.

Uncurable vision loss can arise from a multitude of causes. Trauma-induced vision loss is common in young people and often results in irreversible blindness^2–4^. The leading causes of blindness and visual impairment are cataracts, retinal degenerative diseases (including glaucoma, age-related macular degeneration (AMD) and diabetic retinopathy (DR))^5^. While cataracts can be cured by surgical interventions, current therapies can only slow down the progression of degenerative diseases. For example, anti-VEGF antibodies, which are established treatments for AMD^6^, show limited efficacy and are associated with significant risk. Vision loss can also arise secondary to systemic diseases, such as diabetes^7^, hypertension^8^, embolic disease^9^, auto-immune diseases^10^ and ischemic diseases^11^. In addition, inherited genetic mutations that cause Retinitis Pigmentosa^12,13^ and Stargardt’s disease^13–15^, are also major cause of retinal degeneration. Current therapies for these retinal degenerative diseases have some efficacy in slowing disease progression, although they do not fully restore vision.

As human retinas lack regenerative capacity^16^, therapies currently being tested in clinical trials or undergoing preclinical validation aim to restore vision. Approaches include cell replacement therapies^17^, retinal prostheses^18^, optogenetics^19^ and gene therapy^20,21^. Despite some promising advances, the efficacy of these treatments remains limited. Whole eye transplantation (WET) might offer an alternative solution. However, while whole organ transplantation has been achieved for the heart, lungs, kidneys, liver, pancreas, and intestines, WET has long been considered medically impossible. Recently, in a first-in-class clinical study, the feasibility of ocular reperfusion was demonstrated^22^, bringing hope that eye transplantation could become a future vision-restoring clinical intervention. However, because the retina, optic nerve and CNS are extremely vulnerable to even temporary ischemia, achieving WET with visual recovery requires development of innovative techniques and approaches that challenge paradigms that have long shaped the eye research field.

In addition to its potential use for WET, maintaining an eye *ex vivo* would also be extremely valuable for pre-clinical testing, which is usually performed using rodents. Because mice retinas lack a macula-like structure, which is the primary site of human retinal degeneration, human or large animal eyes are needed. When the retina is isolated from the eyecup and maintained in physiologically balanced oxygenated medium, light-responses can be detected^23–26^. Although, in the absence of circulation, the human retina rapidly loses its capacity for light response^27^. To date, it remains unknown whether retinal function can be preserved in intact eyes *ex vivo* rather than in isolated retinas, as previously reported^25^. Restoring retinal function through the eye’s native circulation is critical for modelling whole-eye physiology and advancing toward transplantation and pre-clinical applications. However, it is a particularly difficult task, given the challenges associated with delivering nutrients and oxygen to the retina inside the post-mortem eye. We hypothesised that if the human eye could be maintained under physiological conditions *ex vivo*, retinal activity could be preserved in the intact eye outside of the body.

To test this hypothesis, we used pig eyes, as they are anatomically similar to human eyes. First, we developed and validated a blood vessel cannulation and perfusion strategy. We then showed that *ex vivo* perfusion of the entire ocular vasculature with an oxygenated solution was sufficient to rescue retinal electrophysiological function for up to 10 hours post-mortem. Furthermore, we also validated our cannulation and perfusion approach using human eyes. Taken together, the ECaBox system preserves retinal structure and viability of the post-mortem eye through efficient perfusion. Consequently, retinal function in the whole eye remains intact, despite a period of ischemia following enucleation. We envision that the ECaBox system, by resuscitating eye globes, could be applied to test retinal therapies *ex vivo*, and potentially for vision-restoring WET.

## Results

### Designing a system for *ex vivo* eye care (ECaBox)

We hypothesised that restoring perfusion to the retinal tissue via physiological circulation could restore function and thus resuscitate the eye. Additionally, we reasoned that to preserve the eye’s functionality, the haemodynamic function in both retinal and choroidal vasculature must be maintained. By optimizing previously reported methods^28,29^, we developed a cannulation and perfusion strategy for pig and human eyes, and incorporated clinical imaging techniques to evaluate the efficacy of retinal perfusion.

To facilitate combining re-perfusion and ocular assessment approach over time, we developed several iterations of an eye support system. We finally obtained a sealable system, the ECaBox (Eyes in a Care Box), with an observation window for optical coherence tomography (OCT) and retinal fundus imaging (Fig. 1a-c, Fig. S1a). A closed-loop perfusion circuit and a control system capable of dynamically regulating both pressure and flow with real-time sensors were also integrated into the system. In addition, ECaBox is equipped with OCT, fundus and fluorescein angiography capability (Fig. S1a).

**Fig. 1.**
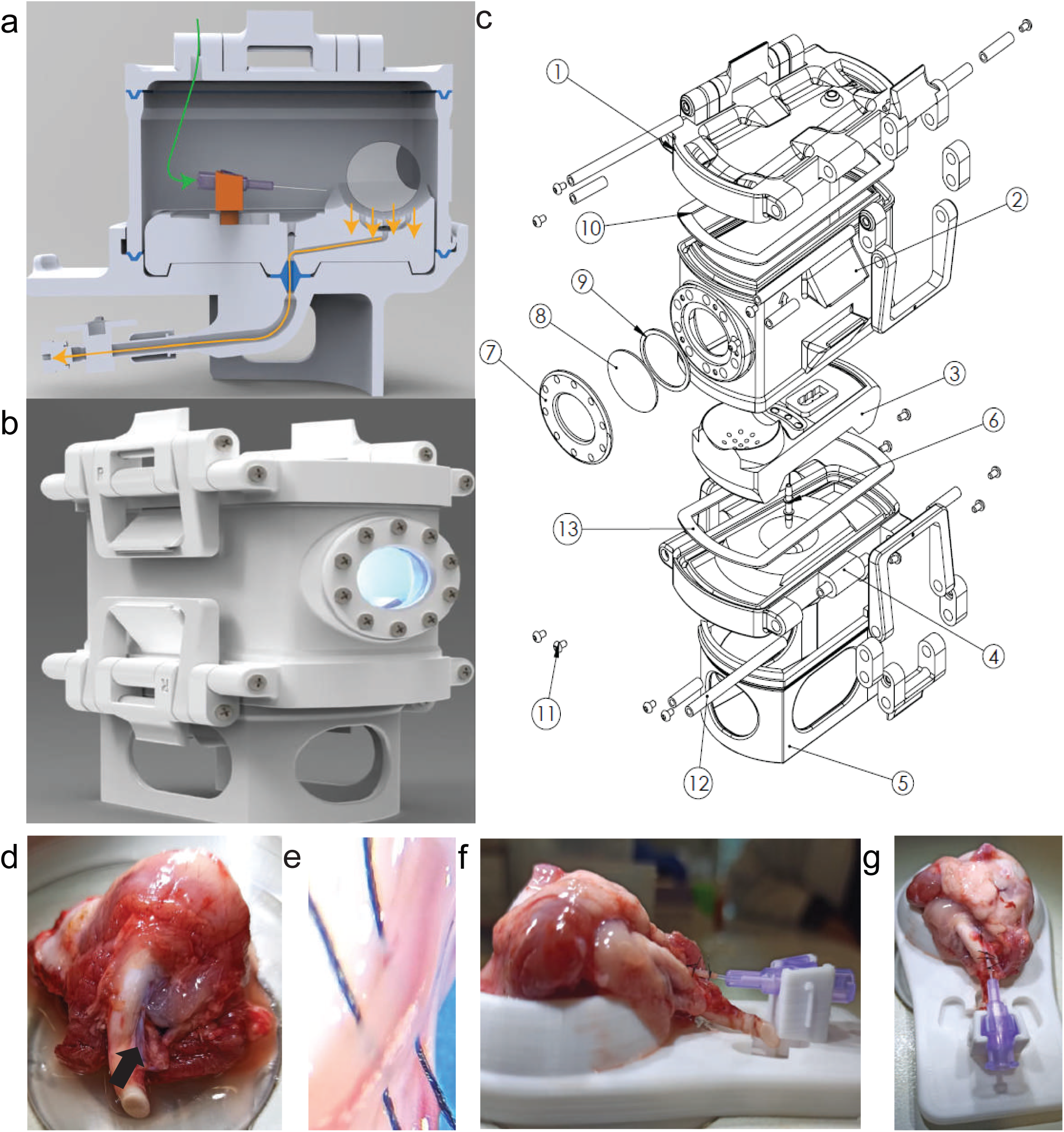
Cannulation strategy and Eye Care Box (ECaBox) design. (a-c) Hermetically sealed ECaBox device. (a) Path of outflowing perfusate (highlighted). The user controls collection of the outflow via the attached 3-way tap. (b) Closing mechanism and OCT window are shown in the anterior portion of the box. (c) Exploded view of the ECaBox system. 1 = Lid, 2 = Main body, 3 = Eye bed, 4 = Base/ drainage system, 5 = Base stand, 6 = Standard Lab Tube Connector, 7 = Window frame, 8 = Window glass, 9 = Window gasket, 10 = Upper gasket, 11 = M3 Screws, 12 = Axis rod, 13 = Bottom Gasket. (d) A pig eye, enucleated and with the muscle resected to make the ophthalmic artery accessible (black arrow). (e) Arterial walls and the mouth of the artery, exposed for cannulation upon further dissection. (f-g) A pig eye placed in the custom-built eye holder with cannula support (26G cannula inserted in the ophthalmic artery), (f) lateral view and (g) overhead view.

The ophthalmic artery was identified soon after the harvesting of pig eyes (Fig. 1d,e), and dissected to facilitate positioning of a 26G cannula (Fig. 1f,g). For the purpose of studying the eye under reperfusion, we developed a supportive holder for the eyeball and a clip-in-cannula support. The holder ensures that the cannula does not move around or fall out of the vessel during manipulation once the eyes are undergoing perfusion (Fig. 1f,g). The imaging window and modular closing components aid in use of the box for various experimental questions (Fig. 1b). The details of all components involved in the set up are shown in an exploded view (Fig. 1c).

### The entire vascular system can be efficiently perfused in ex vivo cannulated eyes

Perfusion with a constant flow rate of 0.5 mL/min, resulted in stable perfusion pressure for over 60 minutes (Fig. S1b). At the same time, it led to a gradual increase in IOP followed by a plateau close to the normal IOP range in pigs, which is around 20-25 mmHg^30,31^(Fig. S1b). To evaluate the efficacy of the ECaBox perfusion strategy, retinal vessel re-perfusion was assessed by retinal fundus imaging and fluorescein angiography performed at room temperature (Fig. 2a and Video S1).

**Fig. 2.**
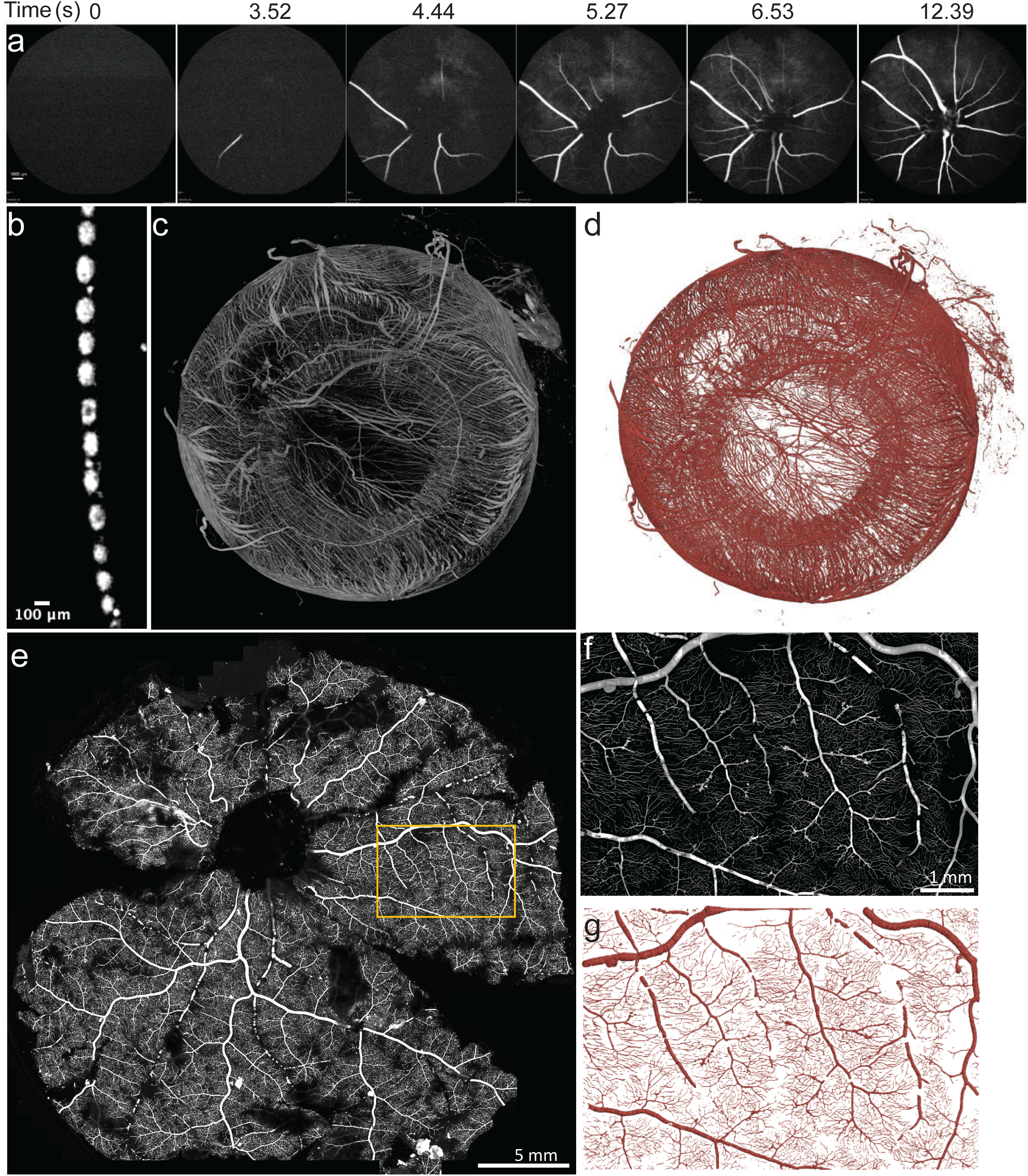
Validation of pig eye perfusion *ex vivo*. (a) Retinal fundus fluorescein angiography (FFA) images of progressive vessel perfusion. Time from the beginning of fluorescein perfusion is indicated in seconds above each image, (*n* ≥ 4) pig eyes. Scale bar 1 mm indicated in the first image is applicable to all subsequent images. (b-c) Representative micro-CT image of the whole eye globe vascular casting, showing perfusion of the choroidal and retinal vascular system. (b) Single-vessel resolution following perfusion and vascular casting. (c) 3h post-mortem, full eye-globe vascular cast, (*n* ≥ 4) pig eyes. (d) AI-segmented and reconstructed ocular vasculature. (e) Confocal max intensity image of the entire retinal z-stack, retinal ganglion cell side up, showing microvascular perfusion after vascular casting. (f) Cropped area from confocal image of the entire retina shown in (e), and its corresponding (g) algorithm reconstructed version of the same area.

Imaging showed arterial and venous flow in approximately 90% of well cannulated eyes (*n* >10) (Fig. 2a, Video S1). OCT imaging showed filling of the retinal vessels on the rostral side of the retina, and the choroidal vessels on the caudal side after 5 minutes of perfusion (Fig. S1c), and retinal edema was absent in the majority of perfused eyes, indicated by no change in retinal thickness after perfusion (Fig. S1d). Having developed the cannulation strategy, we conducted vascular casting to map the vessels within the entire *ex vivo* eyes (Fig. 2b-g). Specifically, the eyes were perfused with BriteVu casting reagent, fixed and imaged by x-ray based Micro-computed tomography (Micro-CT) scanning (Fig. 2b,c).

Using Micro-CT, individual perfused vessels were resolvable (Fig. 2b), and the entire ocular vasculature, including the retina, choroid and iris, was efficiently filled with the casting material (Fig. 2c, Video S2). The vessels could be segmented and reconstructed using the Attention U-Net^32,33^ supervised deep learning model (Fig. 2d,Methods). Vessels were segmented to identify centre lines and diameters, and this initial tree topology was further improved via post-processing (which removed spurious connections and re-joined interrupted segments) (Fig. S2). Specifically, small cyclical vessels were identified (yellow) and further analysed to find processing artefacts (red) for removal. Then, nearby terminal segments were matched for restoration of connection (purple), which, together with the unmodified segments (green) constituted the final output. Thereby, a pipeline for ocular vessel reconstruction was established.

The retina was further examined using confocal microscopy to confirm whether the smaller capillaries below the resolution of Micro-CT were also perfused (Fig. 2e). Casting material was clearly visible in retinal micro-vessels, evidencing their efficient perfusion. A magnified segment of the confocal retinal image reveals perfusion down to the capillary level (Fig. 2f). The clarity of capillary perfusion is further evidenced by automated segmentation (Fig. 2g).

Accessing and extracting a human eye while preserving an intact ophthalmic artery and adhering to ethical limitations, is logistically challenging. Nevertheless, we developed a human eye enucleation approach by adapting a previous method^34^ and the one we developed for porcine eyes. After cannulating and perfusing human donor eyes (Fig. S3a,b), we successfully carried out retinal fundus imaging, showing vascular perfusion with white casting material (Fig. S3c). Furthermore, we visualised the white casting material in the retinal vessels by OCT, suggesting efficient perfusion of human retina (Fig. S3d).

### AI-based reconstruction of vascular anatomy and quantification

Although global vascular casting was successfully performed across multiple samples and at different post-mortem time points, we observed some variation in the extent of penetration. To quantitatively compare these differences across eyes perfused at different post-mortem times, we generated a detailed reconstruction of the vascular anatomy using Attention U-Net model (Fig. 3a and Table S1). The eyes were analysed globally, as well as regionally (by octants – Fig. 3b), and according to the vessel radius groupings. The samples were also designated into early (≤5 hours) or late (≥10 hours) groups according to the time of casting post-mortem.

**Fig. 3.**
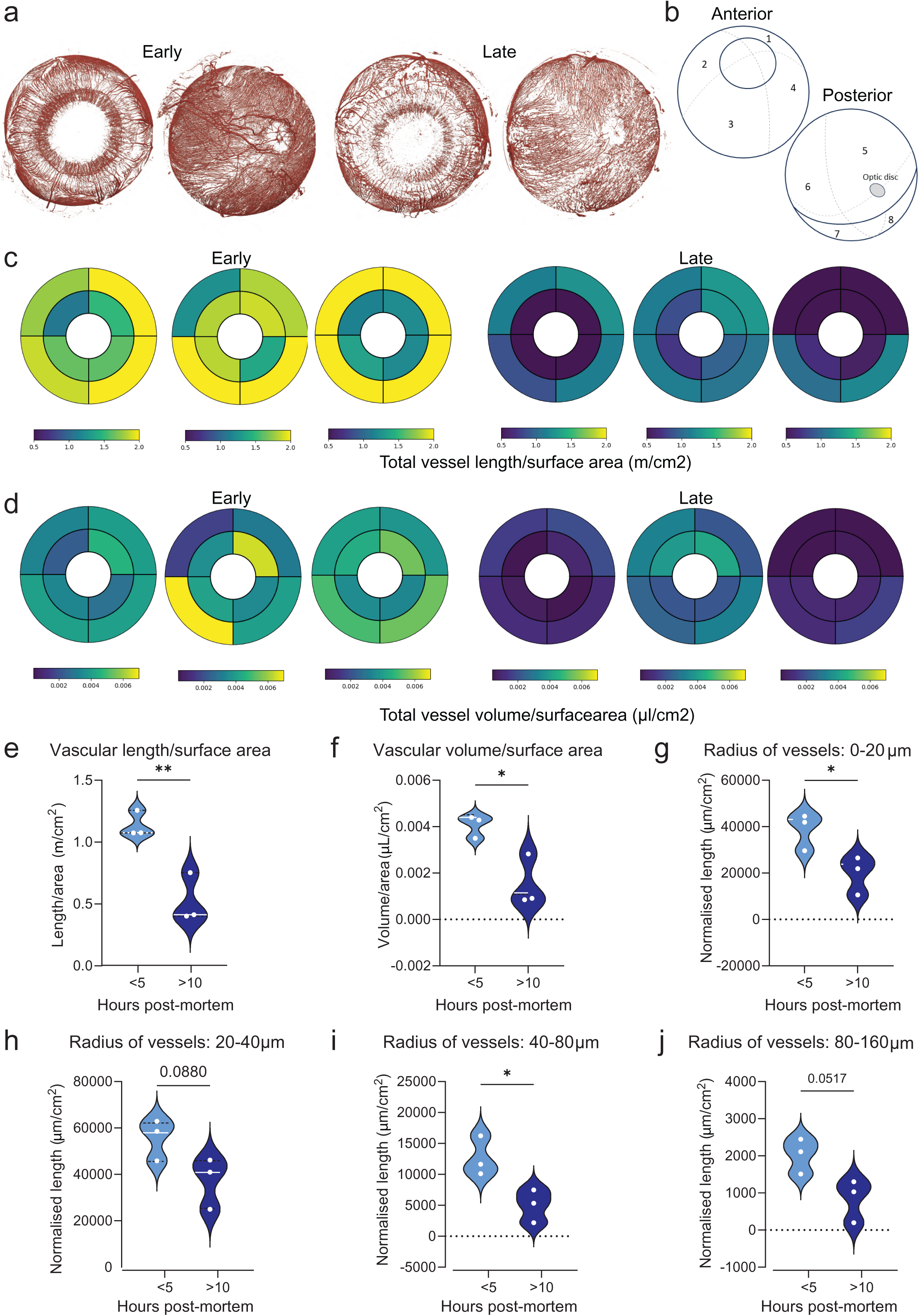
**Vascular parameters in post-mortem eyes at early (≤5 hours) and late (≥10 hours) timepoints**. (a) Sample of full vascular geometries in the anterior and posterior chambers of early and late post-mortem pig eyes. (b) Schematic of the partition of the globe into octants and bulls-eye representation. (c) Total vessel lengths and (d) vascular volumes expressed as density over octant surface area for samples in early and late groups, (*n* = 3) pig eyes per group. Each plot represents one eye. (e-j) Early timepoint (<5 hours) shown in celeste violin plot, late timepoint (>10h) shown in navy. (e) Total vessel length normalised by surface area. (f) Total octant vessel volume normalised by surface area. (g-j) Total vessel length density for different vessel radii groups normalised to surface area: (g) 0-20, (h) 20-40, (i) 40-80, (j) 80-160 µm. Data are compared by student’s t-test *p≤0.05, **p≤0.01.

In each of the samples, the total length (Fig. 3c) and volume (Fig. 3d) of the vascular segments contained within each octant were quantified, and then normalized by the octant surface area to account for possible differences in globe size. This analysis suggested a higher level of vascular density in the early group.

Total vascular length density was lower in the late group compared to the early group, when considered globally (Fig. 3e). Similar reductions in the later post-mortem time were obtained for total vascular volume (Fig. 3f). Vessel lengths could be disaggregated by vessel radius to assess vessels of different sizes. The late group had shorter vessels than the early post-mortem group in the 0-20um and 40-80um radius classes, with a similar trend observed in the other radius classes (Fig. 3g-j). These data indicate that ocular vessels of eyes perfused at later post-mortem times (>5 hours) may lose integrity over time post-mortem, compared to those perfused at early post-mortem times (≤5 hours). When octants were individually examined for length density (Fig. S4a) and volume density (Fig. S4b), all areas of the eyes tended towards a reduction in length and volume in the late group, with most but not all areas showing statistically significant reductions.

### Retinal structure in post-mortem non-perfused eyes

To identify the strategy that best preserved retinal viability, we first characterized retinal degeneration in intact *ex vivo* eyes over time by histologically analysing retinal structure (Fig. 4). We reasoned that at least 24 degrees Celsius (^◦^C) would be required for assessing pre-clinical therapies in *ex vivo* eyes, particularly where electrophysiology is used to evaluate function^35^. Moreover, a previous report found that the retinal light-driven activity in explanted retinas was extended when maintained at room temperature (24^◦^C) compared to 37^⁰^C^36^.

**Fig. 4.**
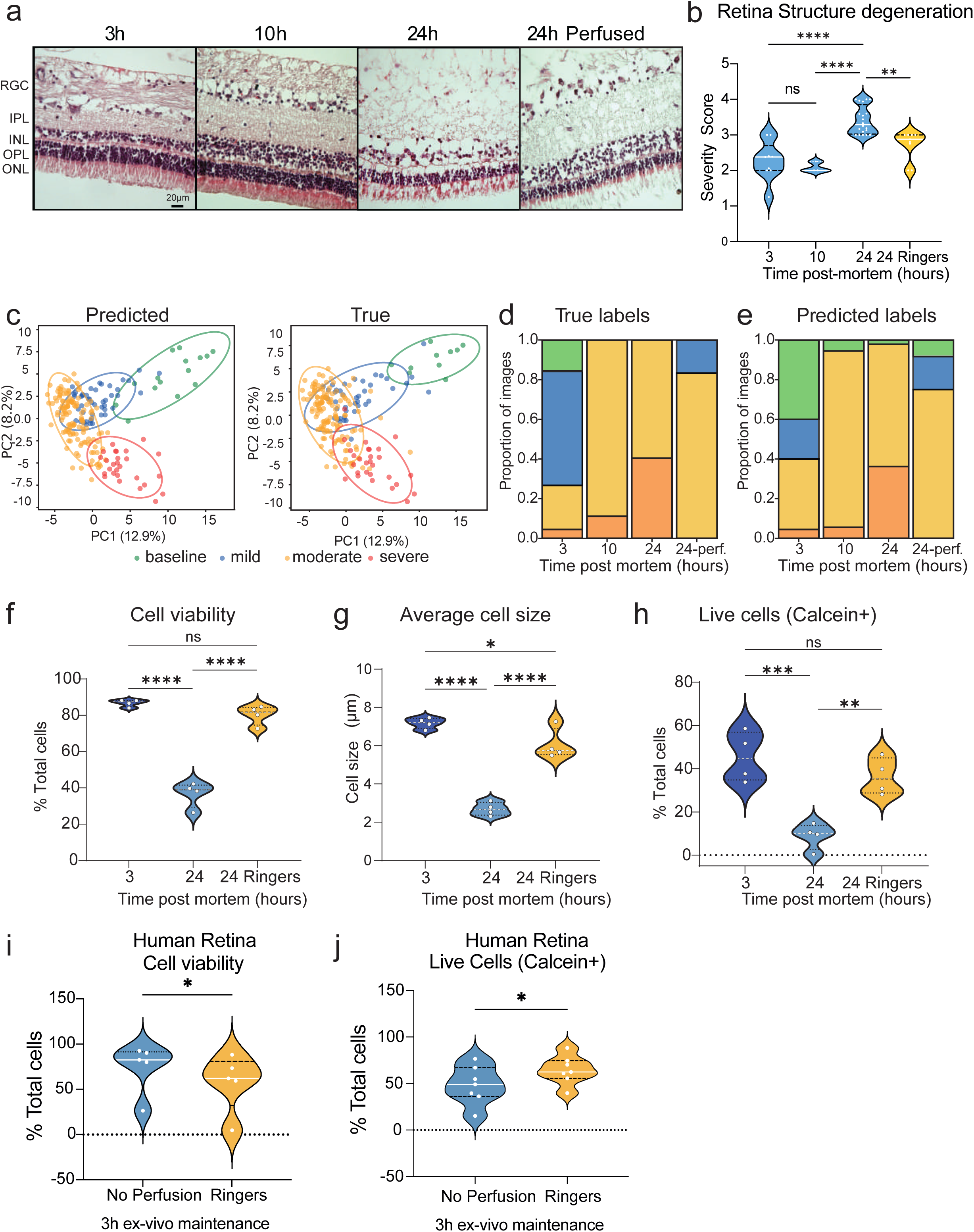
Retinal structure and cell viability in the first 24 hours post-mortem. (a) Representative images of the central retina after maintenance for 3-, 10- or 24-hours post-mortem without perfusion, or 24 hours with Ringer’s solution perfusion at 0.5 mL/min, at room temperature (24 ⁰C). (b) Grading of retinal degeneration of H+E-stained retinal sections from eyes stored for 3-, 10- or 24-hours post-mortem. Retinas were examined in the central, mid and peripheral zones. Post-mortem intervals are indicated on the x-axes, and the violin plot colours indicate no perfusion (blue) and Ringer’s perfusion (yellow). The white points inside each violin plot indicate one pig eye, (3, 24, 24h Ringers, n = 9-20 pig eyes per group, for 10h, *n* = 3). (c) PCA of EfficientNet-B0 features showing true labels and predicted labels in the four severity clusters. (d) Distribution of true labels of the best fold test dataset used to evaluate the performance of the trained model. The groups of unperfused 3h, 10h and 24h post-mortem and perfused 24h post-mortem eyes are comprised by 45, 18, 47 and 12 images, respectively. Only the images corresponding to the central region of the retina are included. (e) Distribution of labels predicted in the trained EfficientNet-B0 model for the same test group. (f-h) Single cells disaggregated from retinas stored in intact pig eyes at 24 ⁰C for various post-mortem intervals were examined for cell viability. (f) Cell counting of disaggregated retinal cells after storage for 3, 24 hours, or 24 hours with Ringer’s perfusion at 500 µL/min, post-mortem. At least 496 cells per eye were counted, over 20,000 cells in the majority of samples. (g) Comparison of cell size alterations over the same timepoints. (h) Flow cytometry analysis of live and dead cells using Calcein as a live cell marker, and normalised to the total number of cells. (f-h) The points inside each violin plot indicate one eye, (*n* = 3-6) pig eyes per group. (i-j) Comparison of cell viability in contralateral eyes from human donors. Cell viability by paired t-test between contralateral eyes after 3 hours perfusion with Ringer’s solution or no perfusion, by (e) countess cell counting (n=5) human eyes and (f) flow cytometry (n=6) for calcein positive live cells. Data with >3 variables are analysed by one Way ANOVA, and contralateral human eyes were compared by paired t-test, ns: P > 0.05, *P ≤ 0.05, **P ≤ 0.01, ***P ≤ 0.001, ****P ≤ 0.0001.

Immediately after enucleation, pig eyes were maintained for 3, 10 or 24 hours at room temperature and then fixed. Retinas isolated from intact eye globes were structurally preserved for up to 10h post-mortem. However, signs of degeneration were present by 24 hours, as evidenced by irregularity and progressive thinning of the nuclear layers (INL, ONL) (Fig. 4a). When we graded retinas for degeneration severity, we did not observe significant differences up to 10 hours post-mortem, while retinas were significantly degenerated by 24 hours (Fig. 4a). Importantly, retinal pigment epithelial (RPE) morphology was well maintained, between early (3 hours) and advanced (24 hours) post-mortem times (Fig. S5), suggesting no clear sign of RPE dysmorphia^37^ at these timepoints.

Retinal degeneration severity was also confirmed by an AI-based scoring method. We trained an EfficientNet-B0 model that accurately classified retinal sections in four different categories: baseline, mild, moderate and severe (Fig. S6a). Feature-space analysis showed four coherent clusters that mirrored both the ground-truth histopathological labels and the network’s predictions (Fig. 4c). Baseline samples (green) occupied a tight, well-defined region at the top of the plot, while the severity classes radiated below in a clear continuum from mild (blue) through moderate (orange) to severe (red), recapitulating the natural progression of retinal degeneration. Confidence ellipses at the 95% confidence level further illustrate that baseline and severe tissues form separate clusters, whereas mild and moderate classes overlap substantially, reflecting the inherent ambiguity in histopathological grading at intermediate stages, consistent with inter-observer variability in clinical practice. Remarkably, the model predictions for 3, 10 and 24 hours post mortem exhibit a similar trend to that of the ground truth, showing an increased shift towards more degenerated states at later post-mortem perfusion times (Fig. 4d,e). Finally, gradient-weighted class activation mapping (Grad-CAM++)^38^ showed highest attention weights of the model within the photoreceptor outer nuclear layer and inner plexiform regions, where degenerative changes are known to occur (Fig. S6b). Activation patterns highlighted an intact laminar architecture in healthy sections, while progressively severe cases showed a concentrated focus on areas of cellular disruption and tissue loss, suggesting that our retinal degradation assessment is robust (Fig. S6b).

Next, we studied the effect of low temperature on tissue preservation. Large organs are commonly preserved at 4^⁰^C, before transplantation to reduce metabolic activity and bacterial proliferation. Although low temperatures are usually inhibitory to retinal function, we examined whether low temperatures preserve structure, protein expression and viability of the retina over post-mortem times. We found that structural degeneration was significantly worse at 24 hours compared to 3 hours post-mortem at both 4^⁰^C and 24^⁰^C preserved eyes, indicating that cold storage is insufficient for preservation of the retinal tissue structure (Fig. S7).

Proteomic analysis showed no changes in overall retinal protein expression between 3-, 10-, and 24-hours post-mortem when stored at 4 ^⁰^C (Fig. S8a). Neither relevant retinal neuronal and glial markers, such as Vimentin, Glial fibrillary acidic protein (GFAP) and Rhodopsin, nor cell-death related proteins changed over these timepoints (Fig. S8b-d). However, both cell size and cell viability were significantly reduced after 24 hours at 4 ^⁰^C, indicating that storage at 4 ^⁰^C is not sufficient to preserve the retina in a post-mortem eye (Fig. S8e-g). Although overall protein expression is stable in post-mortem tissue within the first day at 4^⁰^C, retinal viability is impaired both at 4^⁰^C and at room temperature 24 hours post-mortem (Fig. S7, S8). Therefore, to resuscitate the eye, an intervention such as reperfusion would be required to avoid damage before 24 hours post-mortem.

### Retinal viability is rescued in post-mortem perfused eyes

We then examined if perfusion with oxygenated Ringer’s solution would be sufficient to resuscitate and keep the retina alive in the post-mortem eye. We maintained retinas from two groups of eyes for 24 hours (one without perfusion and one under perfusion with Ringer’s solution) and compared these with eyes analysed 3 hours post-mortem without perfusion (Fig. 4). Retinal structural degeneration severity was greatly reduced in Ringer’s-perfused eyes 24 hours post-mortem compared to non-perfused eyes (Fig. 4a,b). This result was also confirmed by predictions made using EfficientNet. Our trained model not only correctly classified increased damage severity as time after death increased, but also the reverted state after perfusion (Fig. 4d). The model predicted comparable proportions of mildly and moderately degenerated images in 10h non-perfused and 24h-perfused eyes (Fig. 4e and Table S2), indicating improvement compared to 24h eyes without perfusion.

Perfusion of eyes with Ringer’s solution was also sufficient to rescue cell viability (Fig. 4f) and retinal cell size (Fig. 4g). The protection of cell viability achieved by perfusion with Ringer’s solution was further confirmed by flow cytometry analysis of Calcein labelled live cells in disaggregated retinas (Fig. 4h). Overall, eyes maintained under perfusion until 24h post-mortem were significantly more viable than eyes maintained for the same time without perfusion (Fig. 4f-h). These data were validated by comparing perfused versus non-perfused contralateral eyes from 6 human donors. We compared retinal cell viability in contralateral eyes from the same human donor, after one was perfused and the other left without intervention for 3 hours after sample obtention. Perfusion rescued retinal cell viability (Fig. 4i), with variability attributable to the heterogeneity of donors (Fig. 4i,j and Table S4).

### Resuscitation of *ex vivo* eyes up to ten hours post-mortem

Next, we tested retinal function following reperfusion of *ex vivo* eyes. We aimed to evaluate whether retinas in perfused eyes respond to light stimulation. Thus, we undertook the challenging task of measuring electroretinograms (ERG) from intact *ex vivo* eyes after light stimulation. Previous reports show that oxygen is a vital parameter for neuronal function^39,40^, and reduced oxygen levels led to lower ERG amplitude^41^. This suggests that blood oxygen is closely related to ERG response, and that the retina is responsive in a wide range of oxygen saturation levels.

To determine whether restoring perfusion of the eye globes could restore the electrophysiological responses of the retina, we perfused *ex vivo* eyes with oxygenated Ringer’s solution, and measured ERG responses over time post-enucleation (Fig. 5a). A positive response was defined by the presence of a characteristic and repeated ERG-waveform with an amplitude higher than 2.5 standard deviations above the noise level (Fig. 5b and Table S3). Negative signals had no repeated waveform (Fig. 5c). Importantly, ERG amplitude increased with light intensity, reaching saturation at the higher intensities (Fig. S9a).

**Fig. 5.**
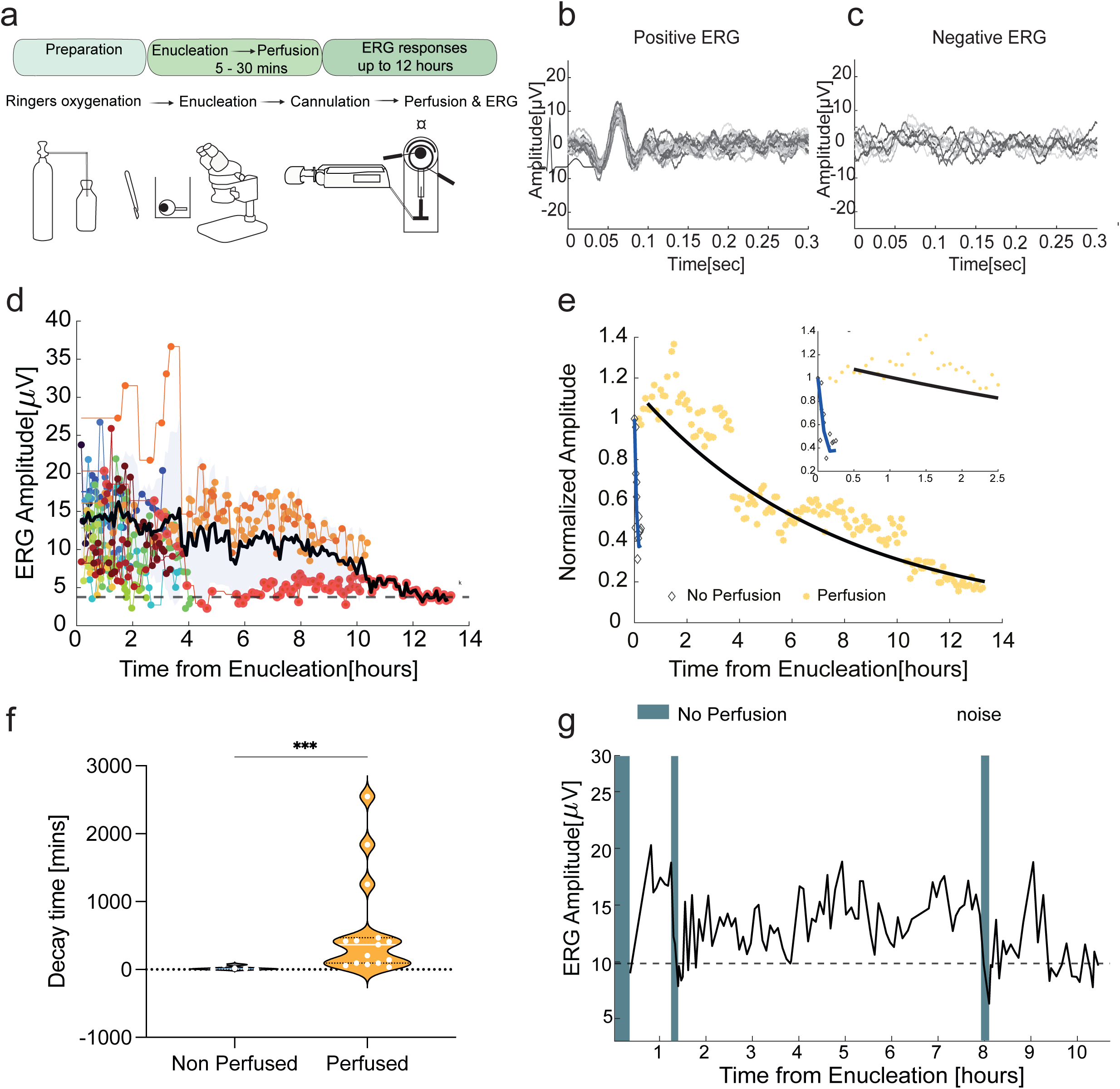
Retinal electrophysiology in perfused eyes over time post-mortem. (a) Schematic illustration of the oxygenation, eye extraction, cannulation, perfusion and ERG, including tools such as forceps, spring scissors and sutures. Cannulation and perfusion are performed within 30 minutes of enucleation. (b) Recording window showing a representative recording from an ERG-positive perfused eye, stimulated at various time points following enucleation, revealed a characteristic signal with stable amplitude (peak-to-peak), for over 3hours post-enucleation. (c) Recording window showing representative waveform where no ERG response is present, at various time points following enucleation in a perfused eye. (d-g) Individual points on each plot represent the ERG amplitude, calculated from the maximum to minimum points of the average waveform, obtained from 30 repeated recordings per timepoint post-enucleation. (d) Overall light responses in whole eyes outside the body, for up to 10 hours of perfusion with Ringer’s solution. ERG amplitudes for all perfused eyes, colours indicating individual eyes, and the average response is shown by the solid black line n =15 pig eyes, noise level indicated by dashed black line. (e) Linear extrapolation of normalized averaged ERG amplitude of all perfused eyes (yellow asterisks) and non-perfused eyes (grey diamonds) and an exponential decay fit of perfused (black line) compared to an average fit of non-perfused eyes (blue line). Insert : zoom in on T=0:2.5 hours from enucleation highlighting significantly faster loss of signals in non-perfused eyes. n = 15 perfused pig eyes, n =5 non perfused pig eyes. (f) Average estimated decay time in minutes in perfused vs non perfused eyes (mean±stdv) n =15 pig eyes. (g) Representative ERG amplitudes for one of three individual eyes whose light responses persisted for at least 10 hours. Light responses are lost when perfusion is switched off (grey-blue background) and revived when perfusion is flowing (white background). n = 3 perfused pig eyes.

ERG signals were detected during perfusion of the eyes with oxygenated Ringer’s solution in 15 out of 36 perfused eyes (41.6% responsive) (Fig. 5d-f and Table S3). ERG amplitude decreased very gradually over the duration of the perfusion (Fig. 5d,e) for up to 10 hours (*n* = 3), and even 12 hours (*n* =1), finally reaching a plateau. In some eyes, recordings were stopped due to logistical reasons rather than the eye ceasing to respond. *Ex vivo* ERG signals were observed in eyes from 15 animals, indicating that the capacity of an *ex vivo* eye to respond is not idiosyncratic to an individual animal’s physiology. The decay of ERG signal due to ischemia during enucleation was immediate in non-perfused eyes (Fig. 5e,f), similar to a previous report^30^. These results highlight the capability of the ECaBox strategy to maintain eye functionality for up to 10 -12 hours after death through perfusion with a readily prepared physiological solution. Notably, within the first 15 minutes after initiating perfusion, no ERG response was observed in any of the eyes. The signal remains close to the noise level (Fig. S9b), which is consistent with gradual resuscitation of activity upon the initiation of perfusion. In three eyes, once a stable ERG signal was established *ex vivo* following perfusion, the perfusion was turned off. As expected, this led to a decrease and eventual cessation of the ERG response. When perfusion was resumed, the ERG signal was restored, further indicating that resuscitation of the retinal activity is dependent on perfusion (Fig. 5g). Future optimization of the entire perfusion system (including perfusate) may result in even longer persistence of the ERG signal.

Finally, we assessed possible causes of the variability of the ERG response detected in *ex vivo* eyes. There was no significant difference in ischemia time, i.e., the time from enucleation to reperfusion, in ERG responsive and unresponsive eyes (Fig. S9c). The baseline *in vivo* ERG amplitude of responsive eyes was compared to the duration of *ex vivo* response, and a very weak positive or no correlation was observed (R squared 0.08865) (Fig. S9d). Moreover, we also compared intraocular pressure (IOP) values obtained in the initial *in vivo* testing between subsequent ex vivo responsive and non-responsive eyes. Here, we found that there was no significant difference, indicating that pre-existing IOP is not the cause of the difference in ERG response (Fig. S9e). ERGs were recorded for all eyes *in vivo* before *ex vivo* recording were performed. There were no significant differences in baseline ERG amplitudes between ex-vivo ERG-responsive and unresponsive eyes (Fig. S9f). These results suggest that additional factors determine *ex vivo* ERG responses and future studies will be required to determine the most pivotal parameters for the successful resuscitation of the retina in an excised eye globe.

## Discussion

The World Health Organization estimates that vision loss affects at least 2.2 billion people worldwide. Therapies that can restore vision are lacking. In this study, we present a unique approach to address this unmet clinical need. Specifically, we leveraged surgical approaches, ophthalmic imaging, viability studies, structural analysis, electrophysiology, and vascular modelling to identify conditions under which the eye can survive outside the body. We present data exploring the *status quo* of the retina when kept in the intact eye following enucleation and a strategy to resuscitate retinal function in the post-mortem eye.

We demonstrate that post-mortem pig eyes can be revived in the ECaBox system. Pig eyes include a “visual streak”, a fovea-like zone responsible for sharp, detailed central vision^42^, and have a similar blood vessel organization and retinal size to human retinas^43,44^. Consequently, the porcine eye in the ECaBox system provides a faithful model of the human eye anatomy, thus facilitating evaluation of treatments in whole eyes *ex vivo* before progressing to live animals and to patients. Furthermore, porcine eyes offer an ethically and sustainably favourable model for pre-clinical testing, as they are similar to human eyes and by-products of the meat industry. Thus, our study supports the 3Rs principle by introducing an alternative approach that reduces the reliance on live animals for experimentation.

Our experimental data demonstrating retinal revival in the intact post-mortem eye represent an important milestone towards achieving vision restoration via WET. Our findings that light stimulation of the retina in intact eyes elicits neuronal responses for multiple hours *ex vivo*, concur with previous studies from isolated retinas^25,45–47^ and retinal organoids^48,49^, and the recent report of light responses in one patient after a side-by-side face allograft and eye reperfusion^22^. Although eye transplant with vision recovery is not currently available to patients, this was a remarkable proof of concept. Angiography showed successful perfusion of the retina, maintenance of eyeball volume, and more importantly, a recorded electrical activity indicating that the visual cortex responds to light stimulation. However, the patient remained unable to see from the transplanted eye, underscoring the vulnerability of retinal tissue.

The novelty of the ECaBox eye maintenance strategy is that it preserves intact eyes for up to 24 hours post-mortem and allows resuscitation of light responses for up to 10 - 12 hours outside of the body, independently of blood. ECaBox allows continuous monitoring of retinal structure and function via real-time imaging and electrophysiological recording in the intact globe. We show that the retina can recover light responses via physiological perfusion through the ophthalmic artery, a response not feasible in eye cup preparations. In the retinal explant system, the electrodes are in direct contact with the retina^29^. In contrast, in our whole-eye system, light responses are detected via an external corneal electrode, as in human patients. The signal intensity recorded from the whole perfused eye outside the body, serves as a clear indication of retinal survival and functional activity. It cannot be directly correlated with *in vivo* or *ex vivo* recordings in retinal explants, as amplitudes are highly dependent on the ERG system set-up, noise isolation conditions, electrode contacts and the biological system – all of which have been adapted here for the unique setting of *ex-vivo* eye examination in a veterinary surgery room.

Thus, while both previous studies^39,40^ and our study support the broader concept that timely oxygenation can restore post-mortem retinal activity, our method establishes proof-of-principle for vascularly mediated revival of the intact eye, representing a key conceptual and technical advance.

Ophthalmic-artery cannulation reported previously^28,29^, were limited to procedural or anatomical characterization of the eye, whereas ECaBox allows continuous monitoring of retinal structure and function as well as of vascular integrity via real-time imaging and electrophysiological recording in the intact globe. Preserving retinal function as long as possible after donation is critical to facilitate WET in a routine clinical setting (where side-by-side donors and recipients is impractical). With future development and refinement of the ECaBox system and a nutrient-rich perfusion fluid, even longer *ex vivo* maintenance of the retina within an intact eye should be achievable.

Human donors show substantial heterogeneity in advanced age and comorbidity, which limits the ability to obtain statistically significant data. Moreover, one of the main reasons for the low number of human donor eyes in our study, is the ethical challenges associated with tissue donation. Currently, we have access only to human tissue that are unsuitable for corneal transplantation, as medical use of donor corneas is prioritized over research use. Now that we have robust evidence demonstrating that we can revive pig eyes, and a solid rationale for attempting the same in human eyes, we are more likely to obtain approval for access to eyes from heart-beating donors, which are typically harvested for transplantation purposes.

Eyes from this type of donors would also allow us to perform ERG experiments on human *ex vivo* eyes. We are planning to develop a portable, surgery-room ECaBox to minimize ischemia in heart-beating donor eyes, when they become available.

In this study we also provided a detailed reconstruction of the entire ocular vasculature, which will enable future work to refine models of vessel architecture and branching connectivity. This will offer the opportunity to further optimize perfusion conditions, by enabling simulated flow analysis on specific samples. A detailed anatomical mapping of the vasculature is a key future component in establishing digital twin capability to model both normal and disease conditions, which will facilitate in silico predictions of therapies, advanced perfusion strategies and pharmacokinetics, before their validation in ECaBox-maintained eyes.

Our data also showed that perfusion preserves retina viability after ischemia and this is in agreement with clinical observations such as those in Amaurosis fugax, a temporary vision loss condition caused by a transient obstruction to the retinal blood flow, in which vision is restored once normal blood flow is re-established^50,51^.

Similarly, temporary reversible vision loss can occur in humans due to conditions such as retinal migraine, retinal vasospasm, central retinal artery occlusion, epileptic seizures and Uhthoff phenomenon. However, upon central retinal artery occlusion, recovery of sight is very limited if perfusion is not restored within 90 min^52,53^. These previous reports suggest that the entire visual system has some degree of tolerance to ischemia, which is in accordance with the revival of light responses reported in the present study.

Finally, our study offers considerations that are important for the preservation of the brain subjected to ischemia. Indeed, while electrophysiological responses are lost upon ischemia, we show that they can be rescued in the neuroretina using an oxygenated solution. This finding may prompt additional reconsideration of clinical protocols and existing paradigms surrounding the utility of re-perfusion in an attempt to reverse brain death.

In conclusion, the ECaBox perfusion and monitoring approaches described in this report demonstrate that retinal light responses can be restored through the eye’s native circulation after death. ECaBox can be used worldwide for evaluation of novel pre-clinical therapies before clinical trials, and for improved preservation of eyes for WET, as it enters clinical practise in the future. Our findings represent a substantial conceptual and technical advance over existing methods, providing a physiologically complete model of post-mortem ocular resuscitation and establishing a foundation for future transplantation and therapeutic research. Ultimately, ECaBox will bring vision-restoring treatments to patients.

## Methods

### ECaBox design, printing and assembly

A supportive device for the eye, including a component which secures the position of the cannula, was designed using computer-aided design (CAD) modelling software, and 3D printed using FormLabs Form3B printer with FormLabs Grey V4 resin (RS-F2-GPGR-04). Laser cut 1 mm silicone gaskets were used to hermetically seal the eye maintenance device.

### Porcine eye obtention from slaughterhouse setting

For retinal structural and viability analyses, porcine eyes were enucleated from recently slaughtered pigs at a local slaughterhouse (Mafrica, Manresa, Spain). The breeds of pigs used in experiments were (Landrace x Pietrain) or (Landrace x Large White) x Duroc. Pigs were stunned into unconsciousness on a carbon dioxide wheel, and suspended by the hind legs before rapid exsanguination, performed by severance of the brachiocephalic trunk at the neck. Exsanguination occurs within 95 seconds, during which time the eyes are extracted. Eyes are extracted by separating the eye globe from the ocular cavity using a paring knife, briefly washed in betadine, and placed in saline with heparin sodium 50 UI/mL, on ice, for transport and delivered to the laboratory within 2.5 hours post-mortem.

### Procedure for obtention of donor consent

Organs and tissue donation were obtained in accordance with Spanish Law 30/1979 on organ extraction and transplantation, and Royal Decree 1723/2012, which regulates the procurement, clinical use, and territorial coordination of human organs. Spanish legislation is based on the principle of “presumed consent,” whereby every individual is considered an organ donor for medical purposes, unless they have explicitly opted out during their lifetime. In this study, we established a specific procedure to confirm this consent for scientific use of tissues, from each donor included in the study. The Donation Coordination Unit of Val d’Hebron hospital (Barcelona) obtained informed consent from the donor’s family members and the individuals directly responsible for the donation after death using a standardized document for each donor included in the study. The reason, purpose, and intended use of the donated ocular tissue were explained in detail to the family members. The donor’s family were also informed about the measures taken to preserve facial integrity, such as the use of appropriate prosthetic devices to replace the eyeballs. If consent for the donation was obtained, this altruistic act was formalized by signing the informed consent form.

### Human eye enucleation

Human donor eyes were collected after explicit written informed consent and approval of the Medical Ethical Committee of the Vall d’Hebron Hospital, Barcelona, Spain, (protocol no. PR(AMI)233/2021 (SO 484/2021)). Donor eyes were used for experiments at CRG under “Parc de Salut Mar Clinical Research Ethics Committee” approval with number 2022/10442/I. Anonymized donor characteristics, including age, sex and primary medical history and cause of death is included in Table S4. The human eye enucleation strategy was developed by experienced Ophthalmologists in collaboration with the donation and transplantation coordination team at Vall d’Hebron Hospital Barcelona, according to a previous method^34^, with some modifications by our group. The medial transconjunctival approach was used. Briefly, a 270-degree conjunctival incision was made 4 mm posterior to the corneoscleral junction. The medial rectus muscle was disinserted from the globe. 4-0 silk traction sutures were passed beneath the superior and inferior rectus muscles. The eye was then pulled to the temporal side and dissection was performed through the orbital fat in a blunt manner. Once the optic nerve was identified, the section was performed as far posteriorly as possible. Finally, the superior, inferior, lateral rectus, and oblique muscles were disinserted. The globe and some of the intraorbital fat were obtained in bloc. After the surgery, a standard local prosthesis was used, resulting in a satisfactory aesthetic result that did not entail facial deformities.

### Cannulation and perfusion of the porcine and human eyes

To prepare porcine eyes for perfusion, the following steps were performed. The conjunctiva was trimmed, and the muscle tissue surrounding the optic nerve was carefully dissected to reveal the ophthalmic artery. This artery lies adjacent to the optic nerve and is distinguishable due to its size and thick arterial walls compared to veins. A 26G cannula (BD Neoflon Ref. 391349) was positioned in the ophthalmic artery and secured using non-absorbable braided silk suture 6/0 (Fine Science Tools Item no. 18020-60). Perfusion was driven by a LongerPump BT100-1L peristaltic, or Braun syringe pump at a rate of 0.5 mL/min. Oxygenation was performed by bubbling solutions with carboxygen (95% O_2_, 5% CO_2_) for 30 minutes prior to use. Human eyes were obtained 6-10 hours post-mortem and were maintained at 4⁰C in saline solution with heparin sodium 50 UI/mL. As previously reported, the feasibility of successfully harvesting the central retinal artery is very dependent on optic nerve^54^. Where >1.5cm optic nerve and >2cm surrounding vascular tissue were collected, and the central retinal artery was present, we performed vascular cannulation as described below for porcine eyes, with adaptation of flow by hand perfusion, based on detected resistance.

### Recording of hemodynamics in porcine eyes

For longer periods of perfusion and hemodynamic measurements, we built a dedicated system consisting of a sterile, closed-loop perfusion circuit that includes a perfusate reservoir (5L tank with 1L submersible disposable reservoirs) with thermal conditioning and oxygenation (LAUDA Alpha RA 8), a peristaltic pump, and a control system capable of dynamically regulating both pressure and flow based on real-time sensor feedback. The device also included a corneal hydration circuit, which delivers temperature-controlled fluid at scheduled intervals. Pressure sensors (MEMSCAP SP854) and flow sensors (Sensirion LD20-2600B) were directly embedded into the circulation line immediately downstream of the peristaltic pump. The flow sensor probes also included temperature sensors in direct contact with the fluid, altogether providing a low-latency, high-fidelity acquisition.

The hardware operation as well as user interface were controlled by a custom software written in LabVIEW 2025 Q1 (National Instruments). PID controllers were implemented and optimized for both pressure- and flow-driven perfusion modes.

### Optical coherence tomography (OCT) and retinal fundus imaging

To evaluate perfusion of post-mortem eyes, the retina was observed using OCT and retinal fundus imaging via the Heidelberg system. Eyes were positioned in the ECaBox eye support, facing the camera. Infrared retinal fundus images were used to observe whether retinal vessels had been filled with white BriteVu casting material. Images were obtained in different focal planes. Where fluorescein was perfused, blue autofluorescence images and videos of vascular filling were acquired. OCT images were acquired around the optic nerve, to understand the depth of retinal penetrance of the casting material. Although in one case, OCT was successfully performed on the fresh human eye, in the case of human donor eyes, where opacities were already present due to the advanced age of donors (Table S4), the casted eyes were fixed, and the anterior chamber was dissected before performance of OCT. This facilitated visualisation of the retina in situ, despite the presence of cataracts and opacities.

### Micro-CT

Micro-CT scanning was performed with a micro-computed tomography Skyscan 1272 (Bruker), with an X-ray tube voltage of 70kV, anode current of 142uA, Al 0.5mm filter, pixel size of 6-12um, rotation step of 0.4°, total rotation of 180° and scan averaging of 5. NRecon (Bruker, version 1.7.0.4) reconstruction software was used to transform raw micro-CT data into tomographic images in 3D in unsigned 16-bit format, with gray level ranging from 0-65535. Volume rendering of the reconstructed images was generated by CTVox (Bruker) to obtain the 3D images.

### Vessel segmentation AI model

Our vascular segmentation pipeline used a supervised deep learning approach. Ground truth data were sourced from 4 micro-CT datasets, which were divided into sub-blocks and 310 were randomly picked to be manually annotated. Training, validation and test sets were assigned at 75:15:10 ratio. 3D U-Net^55^, V-Net^56^, Deep Residual U-Net^57^ and Attention U-Net^32,33^ were trained 5 times each and top F1 scores were compared. As shown in Table S1 (best results shown by bold lettering), Attention U-Net was most performant and was used for the subsequent processing. The segmented vascular mask image was skeletonized and converted to a sparse graph format yielding an effective data compression from around 70GB to a few kilobytes. Vessel diameters were detected using a modified version of Rayburst sampling^58^, applied in a predictor-corrector fashion, which initially used a local thickness filter as a first pass to tune a truncated, oriented sampling core that is adaptively scaled by the thickness. Full width half maximum criterion was used for ray termination.

Connected components were identified from the full graph, which often revealed smaller subtrees isolated from the main network. The detachments were assumed to have been introduced post-casting (since occlusions would not have allowed Britevu to penetrate distally), and thus subtrees were included in the quantification. In addition, topological irregularities such as spurious connections between parallel vessels and minor gaps were corrected (Fig. S2). To do this, first, we searched for all simple cycles of bounded length in the graph, and then detected and removed problematic connections using heuristic criteria based on diameter fluctuations along the vessel segments. Following this, terminal segment search and direction matching were used to join up short gaps between segments which were present due to contrast fluctuations.

### Evaluation of Regional Vessel Networks

To enable a comparison between different casted eye globes, all reconstructed vascular network meshes were aligned to a standard orientation using the iris plane (from which anterior-posterior axis was identified), and the optic disc centre (which was selected to be the 0 angular position). The equator of the globe, which was assumed to bisect the front and rear hemispheres (Fig. 3a), was then defined, finally resulting in the division of the globe into 8 parts (Fig. 3b). The vascular segments contained within each octant were identified, and the total vascular length and volume, and corresponding densities were calculated.

Paraffin processing of dissected retinae

For preparation of paraffin embedded retinal sections, the anterior chamber of the eye, including the lens and vitreous humour, were removed and the remaining posterior chamber eyecup was fixed in 4% PFA at 4 ⁰C overnight. The following day, retinas were dissected out of eyecups, and wax processed and embedded for paraffin section staining. Paraffin sections were cut at a 5 µm thickness and mounted on super-frost slides. Haematoxylin and Eosin staining was performed on paraffin-embedded 5µm thick retinal sections, from the central, mid and peripheral retinas. Histology images were captured using a Leica DM6000B microscope.

### Immunostaining of RPE flatmounts

Following overnight fixation of the eye cup in 4% paraformaldehyde (PFA), RPE-choroids were dissected away from the sclera and retina. RPE flatmounts were washed with phosphate buffered saline (PBS) and blocked with 10% goat serum for one hour at room temperature. Alexa Fluor™ 488 Phalloidin (# A12379, Thermofisher) was applied at a dilution of 1:1000 in 5% goat serum in PBS, for 2 hours at room temperature. Flatmounts were washed 3 times in PBS, and mounted with VECTASHIELD® Antifade Mounting Medium (H-1000-10, Vector laboratories) in 50mm FluoroDish glass bottomed dishes (FD5040-100, World Precision Instruments). Flatmounts were imaged using a Leica CTR7000 HS microscope, and RPE cell structure was analysed using Cellpose.

### Analysis of RPE cell morphology

Phalloidin labelled images of the RPE cells of porcine retinal explants were analysed using Cellpose. Images were denoised and segmented using a diameter of 200 pixels and a flow threshold of 0.2. A mask of the segmentation outline of the Phalloidin positive areas was created, representing the borders of the RPE cells. RPE cell shape descriptors, namely cell area and cell shape were obtained. The average (most commonly occurring area) was compared between eyes maintained for different durations post-mortem.

### Histology grading

Histology sections were graded according to the severity scale outlined previously^62^ by 2 blinded evaluators. Briefly, damage to the inner and outer nuclear layers was classified according to the following criteria. Retinas were deemed healthy when the typical retinal layers (ONL, OPL, INL, IPL, GCL), were present, without any damage. The presence of any tears in either the ONL or INL meant an image had at least a low severity of damage. Loss of nuclear density in some areas increased the severity to moderate degeneration, and uniform loss of nuclear density was graded as severe degeneration.

### AI-based classification of the retinal structural degeneration in histology sections and interpretable AI analysis

To support our classification of retina images in four different states of structural degeneration (baseline, mild, moderate and severe) we followed previous approaches that use deep learning to solve classification tasks^59^. We implemented RPD-Net, the convolutional neural network (CNN) with the architecture demonstrating best classification accuracy in the referenced publication, and EfficientNet-B0, a model commonly used for classification tasks^60^. In both cases, we used the evaluations performed by a group of observers to train the model to reproduce the assessment by itself and performed 5-fold cross-validation. Finally, we calculated the weighted average of F1 score across all classes, achieving a mean F1 score of 0.84 in the best case of EfficientNet-B0 and 0.46 for the best RPD-Net fold. According to these results, we decided to use EfficientNet-B0 as out best-performing model for image classification.

The models were trained using a dataset comprising 734 histological images obtained from 44 different pig eyes collected at 3-, 10-, and 24-hours post-mortem, representing four distinct degeneration states (34 baseline, 141 mild, 467 moderate, and 92 severe). Both perfused and non-perfused retinas were included, but perfusion state was not provided as an explicit label to the model to maintain unbiased classification. We performed 5-fold cross-validation, generating five random independent test sets of 221 images each. For each training split, the remaining 513 images were augmented using random crops and rotations to expand the dataset to 15,903 images, of which 12,723 were used for training and 3,180 for validation. Model performance was evaluated using accuracy, F1 score, and confusion matrices averaged across the five folds.

To analyse model representational features, we extracted 1280-dimensional embeddings from the global average pooling layer of EfficientNet-B0 and projected them onto two principal components using principal component analysis (PCA). The proportion of explained variance and class covariance structure were visualized, and 95% confidence ellipses were computed to assess cluster dispersion (Fig. 4c). To confirm that the model’s decisions were based on histologically relevant features, we applied gradient-weighted class activation mapping (Grad-CAM++). Activation maps were generated for images across all severity classes to identify the regions contributing most strongly to classification (Fig. S6b).

### Flow cytometry

For flow cytometry experiments, retinas were isolated from intact eye globes after maintenance for 3, 10 or 24 hours post-mortem at 4 ⁰C or at room temperature in saline with heparin sodium 50 UI/mL. For analysis of live retinal cells, retinas were disassociated in 800 µL Papain containing DNAse in a shaker at 650 RPM at 37 ⁰C, for 30 minutes. After digestion, the cell suspension was spun down at 200 RPM at room temperature for 5 minutes to pellet the cells. The digested cells of one entire retina were resuspended in 2 mL PBS. For staining, 100 µL of the retina cell suspension was added to 1900 µL PBS and 2 µL Calcein-AM was added. Calcein-AM staining was conducted for 20 minutes on ice before cytometry. Dyes were not washed before cytometry. Immediately before cytometry, 2 µL DAPI was added to each sample. Samples were filtered using Falcon® 5 mL Round Bottom Polystyrene Test Tube, with Cell Strainer Snap Cap, product no. 352235. Flow cytometry was performed using a LSRII cytometer and analyzed using FlowJo. The population of cells were identified from all events detected, using Compensated IndoViolet A versus FSC-A gates. Events below 50K FSC-A were considered debris, and the events >50K FSC-A were considered cells. The calcein positive live cells population was identified and represented as a percentage of total cells.

### Porcine eye obtention and perfusion in surgical setting

For ERG experiments, *in vivo* ERG was performed on anaesthetised, intubated pigs, of both sexes, 35-52 kg in weight, in a veterinary surgery at Universitat Autonoma de Barcelona. Animal handling was conducted in accordance with the European Council Directive 2010/63/EU and the Spanish legislation (RD53/2013).

ERG was performed on terminally anaesthetised animals under deep, terminal general anaesthesia to ensure the complete abolition of consciousness and pain perception and the animals did not regain consciousness at any point. The animals were anesthetized using the following protocol. Premedication (sedation + analgesia) consisted of azaperone 4 mg/kg (Sediron®, Livisto SL, Spain) + ketamine 10 mg/kg (Anesketin®, Eurovet Animal Health B.V., Netherlands) + midazolam 0.2 mg/kg (Dormazolam®, Dechra Regulatory B.V., Netherlands) + methadone 0.2 mg/kg (Insistor®, VetViva Richter GmbH, Austria) administered intramuscularly (IM). After 15 minutes, a catheter was placed in the auricular vein, the animals were preoxygenated with 100% oxygen, and anesthesia was induced with isoflurane (Isovet®, B. Braun VetCare SA, Spain) at 4–5% using a face mask. Once the animals had lost the swallowing reflex, endotracheal intubation was performed, and anesthesia was maintained with isoflurane at 1.5–2% via a circular system with 100% oxygen.

An experienced porcine veterinarian was in charge of all experiments, ensuring welfare management by personnel competent in species-specific handling, anaesthesia, monitoring, and euthanasia techniques, further minimising any risk of suffering. The protocol was designed by the veterinarian to prevent pain or distress at all stages, in alignment with the 3Rs principles, particularly refinement. Key physiological indicators—including heart rate, respiratory rate, reflexes, and oxygenation—were used to assess the depth of anaesthesia throughout the procedure.

At the end of the procedure, euthanasia was performed while the animals were still under anaesthesia, using an approved and humane method (200 mg/kg of pentobarbital), ensuring a painless and distress-free death.

Once *in vivo* functional ERG was confirmed, enucleation was performed. Eyelids were secured open using a blepherostat and grasped using 3 Pean Forceps. Exenteration was performed using an 11-blade scalpel and mild traction on the third eyelid using the Pean Forceps. The eyes and surrounding content were placed immediately in saline solution (to wash off excess blood), and moved to a dark room where the eye was covered in aluminium foil to facilitate light adaptation while it was cannulated. The ophthalmic arteries were immediately cannulated and stabilised in a custom-built eye holder and cannula support, where they were kept for the duration of ERG recordings. Once the cannula entered the artery, the artery was washed with Ringers solution by hand perfusing 1-2 mL. The cannula was secured using 4.0 braided sutures. Perfusion was commenced immediately, at a rate of 0.5 mL/min, using a B. BRAUN Perfuser Space Syringe Pump, no.8713030. Once cannulated, eyes are perfused with freshly prepared carboxygenated Ringer’s solution (110 mM NaCl, 22 mM NaHCO_3_, 2.5mM KCl, 1.6mM MgCl, 1mM CaCl_2_, NaH_2_PO_4_, and 10mM) at a flow rate of 30mL/hour and kept at room temperature, around 24°C.

### Electroretinography (ERG)

Two phases of ERG examinations were performed per eye: *in vivo* recordings and ex vivo recordings. In the live anaesthetised animal, pupils were dilated using compound tropicamide eye drops (5 mg/ml) and eyes were hydrated electrode contact gel (Parker laboratories) used to avoid corneal dehydration and improve the signal transmission. Animals were dark-adapted for at least 15 minutes before recordings. For both *in vivo* and *ex vivo* recordings, all recordings were performed using a custom-made ERG system, consisting of electrooculogram amplifier (BioPac, Ref. EOG100C), isolated power supply module (BioPac, Ref.IPS100C), Multifunctional data acquisition device (National instruments Ref. USB-6001), disposable corneal electrode ERG-Jet (Fabrinal Ref. F-06), disposable subdermal needle electrode (Technomed Ref. TE/S43-438) and a blue LED used as flash stimulus controlled by a purpose-built software interface, built in MatLab. Both *in vivo* and *ex vivo* eyes were stimulated with light power of 500nW/mm^2^ (on the cornea), with 20 msec pulse duration, 1 sec rest and a total of 30 pulses. The recorded data were then analysed.

### ERG data analysis

Firstly, the 50Hz noise was filtered using readily available MATLAB functions, followed by the bandpass filtering of the data (1-100Hz) to extract the ERG signal. The average of the 30 repetitions was calculated and the ERG amplitudes were then defined as the difference between the maximal and minimal values of the average signal. The noise level was defined as 2*STDEV of the average signal amplitude of the time window where no response is expected (400-900ms from stimulus onset). *In vivo* and *ex vivo*, noise levels were determined for each eye after electrode placement, by conducting a stimulus with the light fully isolated in aluminium foil or with the trigger activated and the light stimulus set to 0, preventing stimulation of the eye. ERG recordings were then performed using the same method described above for the *in vivo* recordings. Eyes are dark adapted within 5 minutes post-mortem, and ERG measurements commence as soon as the perfusion cannula is secured (usually 15-30 minutes post-mortem). ERG signals were then recorded approximately every 5 minutes, until the signal was indistinguishable from the noise.

### Intraocular pressure recordings

Intraocular pressure in *in vivo* pigs and *ex vivo* eyes was measured using TonoVet handheld tonometer. Ocular moisture was maintained during IOP readings using HYLO GEL® lubricating eyedrops drops.

### Proteomics sample preparation

Retinas were extracted from post-mortem porcine eyes stored at 4 degrees, at 3, 10 and 24 hours post-mortem. Protein was extracted with RIPA (NaCl 150 mM, 1% IGEPAL® CA-630, 0.5% Sodium Deoxycholate, 0.1% SDS, Tris 50 mM, pH 8.0.) + protease inhibitors. 10 µg of each sample were digested with trypsin and endoproteinase Lys-C.

Samples (10 µg) were reduced with dithiothreitol (30 nmol, 37 °C, 60 min) and alkylated in the dark with iodoacetamide (60 nmol, 25 °C, 30 min). The resulting protein extract was first diluted to 2M urea with 200 mM ammonium bicarbonate for digestion with endoproteinase LysC (1:10 w:w, 37°C, over 6h, Wako, cat # 129-02541), and then diluted 2-fold with 200 mM ammonium bicarbonate for trypsin digestion (1:10 w:w, 37°C, over 8h, Promega cat # V5113).

After digestion, peptide mix was acidified with formic acid and desalted with a MicroSpin C18 column (The Nest Group, Inc) prior to LC-MS/MS analysis.

### Chromatographic and mass spectrometric analysis

Samples were analyzed using an Orbitrap Eclipse mass spectrometer (Thermo Fisher Scientific) coupled to an EASY-nLC 1200 (Thermo Fisher Scientific). Peptides were loaded directly onto the analytical column and were separated by reversed-phase chromatography using a 50-cm column with an inner diameter of 75 μm, packed with 2 μm C18 particles (Thermo Fisher Scientific, cat # ES903).

Chromatographic gradients started at 95% buffer A and 5% buffer B with a flow rate of 300 nl/min and and gradually increased to 25% buffer B and 75% A in 105 min and then to 40% buffer B and 60% A in 15 min. After each analysis, the column was washed for 10 min with 100% buffer B. Buffer A: 0.1% formic acid in water. Buffer B: 0.1% formic acid in 80% acetonitrile.

The mass spectrometer was operated in positive ionization mode with nanospray voltage set at 2.4 kV and source temperature at 305°C. The instrument was operated in data-independent acquisition mode, with a full MS scans over a mass range of m/z 500-900 with detection in the Orbitrap at a resolution of 120,000. The auto gain control (AGC) was set to 1e6 and a maximum injection time of 246ms was used. In each cycle of data-independent acquisition analysis, following each survey scan, 40 consecutive windows of 10 Da each were used to isolate and fragment all precursor ions from 500 to 900 m/z. A normalized collision energy of 28% was used for higher-energy collisional dissociation (HCD) fragmentation. MS2 scan range was set from 350 to 1850 m/z with detection in the Orbitrap at a resolution of 30,000. The AGC was set to 1E6 and a maximum injection time of 54 ms was used.

Digested bovine serum albumin was analyzed between each sample to avoid sample carryover and to assure stability of the instrument and QCloud has been used to control instrument longitudinal performance during the project.

### Data Analysis

Acquired spectra were analyzed using a library-free strategy with DIA-NN v1.8.1 software^61^. The data were searched against the Sus scrofa database UP000008227. For peptide identification trypsin was chosen as enzyme and up to one missed cleavages were allowed. Oxidation of methionine was used as variable modification whereas carbamidomethylation on cysteines was set as a fixed modification. False discovery rate (FDR) was set to a maximum of 1% at both peptide and protein level. Precursor and fragment ion m/z mass range were adjusted to 500-900 and 350-1850, respectively. For peptide quantification match-between-runs was enabled, protein inference was set to ‘Protein names (from FASTA)’ with ‘relaxed-prot-inf’/’Heuristic protein inference’ option and the quantification strategy was set to ‘Robust LC (high precision)’.

MSstats v4.10, an R based package tool designed for the statistical analysis of quantitative proteomic data^62^, was used to perform a paired statistical inference between the three different time points present (T03H, T10H and T24H). The raw proteomics data have been deposited to the PRIDE repository with the dataset identifier PXD073299.

### Statistical analysis

Throughout the manuscript, each point on the plots represents one eye. In all figures, only one eye per animal were included, due to constraints at the slaughterhouse, except in Fig. 5 and Fig. S3, where both eyes from the animals and human donors were accessible and included. Normally distributed data were compared by student’s t-test, One-Way ANOVA or Two-Way ANOVA with multiple comparisons tests, according to the number of variables assessed.

## Additional Information

AI vascular segmentation networks were implemented using open-source package MONAI^63^. Topology analyses were performed with the open-source library networkx 3.4.2^64^. RPE cell segmentation was performed using Cellpose : a generalist algorithm for cellular segmentation^65^.

EfficientNet-B0 model training and inference for retinal degeneration classification were implemented using PyTorch (version 2.8.0) and torchvision (pretrained ImageNet weights). Principal component analysis and feature-space visualisation were performed using scikit-learn (v1.7.1) and matplotlib (v3.10.3). Grad-CAM visualisation for model interpretability was generated using the PyTorch library for CAM methods^66^.

Any additional information required to re-analyze the data reported in this paper is available from the lead contact upon request.

## Acknowledgements

This work was supported by: Future and Emerging Technologies (FET) grant from the European Commission, Grant agreement ID: 964342 to M.P.C., Y.M and J.L; Ministerio de Ciencia e Innovación [PID2020- 114080GB-I00 / AEI / 10.13039 / 501100011033, BFU2017- 86760-P (AEI / FEDER, UE) to M.P.C.; AGAUR grant from Secretaria d’Universitats i Recerca del Departament d’Empresa i Coneixement de la Generalitat de Catalunya [2017 SGR 689 and 2021-SGR2021-01300] to M.P.C.; We acknowledge support of the Spanish Ministry of Science and Innovation through the Centro de Excelencia Severo Ochoa (CEX2020-001049-S, MCIN/AEI/10.13039/501100011033), and the Generalitat de Catalunya through the CERCA programme.

Authors would like to acknowledge: Aida Rebollo Morell for accompanying us in initial efforts to achieve perfusion of porcine eyes; Carlotta Viana for aiding in aesthetic improvement of the figures; Veterinarian Xavier Moll Sanchez at Universitat Autónoma de Barcelona for experimental assistance and advice; Diego Gutierrez’ 3D printing design services; Universitat Polytecnica de Catalunya’s Micro-CT services; Mafrica, Manresa, Spain, for the provision of porcine tissues; Nicky van Kronenburg, Charis Rousou, Enrico Mastrobattista at Utrecht University, for advice on porcine eye extraction and cannulation; Justin D’Antin and the research department at Barraquer Institute for Ophthalmology, for their interesting discussions and suggestions; We are grateful to UPF/CRG Flow Cytometry Unit, CRG Proteomics Unit, CRG Advanced Light Microscopy Unit for their support and assistance with this work.

## Author contributions

M.P.C conceived and supervised the project; E.M.B designed and conducted experiments in consultation with U.D.V, M.C.A, N.F, J.L, Y.M and M.P.C; E.M.B, R.C.M, M.C.A and U.D.V developed porcine eye cannulation protocol; A.S co-ordinated informed consent obtention and donor logistics at Vall d’Hebron hospital; R.C.M, J.A.M.F and J.J.P.G performed human eye enucleation; E.M.B, M.C.A and D.C.R performed human eye perfusion, viability assessment and imaging; R.C.M and Y.M provided ophthalmology consultancy to the project; E.M.B performed porcine eye perfusion evaluation experiments, flow cytometry, cell viability, immunohistochemistry and related analysis; E.M.B, U.D.V, N.F performed electrophysiology experiments; N.F performed electrophysiology data analysis, assembled the system and wrote the system control software; M.S.I and J.L performed segmentation of vascular casting and analysis of vascular segmentation; M.C.A provided technical support in various experiments, and prepared samples for proteomics; M.F.M conducted cellpose analysis on RPE flatmounts; M.F.M and M.R. conducted AI analysis of histology sections; M.G.S assisted with acquisition of histology images; E.M.B and D.C.R performed IOP measurements, OCT and pressure-control perfusion experiments, M.B and J.L wrote the software and performed analysis; E.M.B prepared the figures, in consultation with M.P.C; R.C.M and A.S.C led the team responsible for human donor eye enucleation; J.L supervised vascular segmentation machine learning and analysis; Y.M. supervised electrophysiology experiments and system design; E.M.B. and M.P.C wrote the manuscript, with input from N.F, J.L, Y.M. All authors discussed the data.

## Supplementary Information

**Fig. S1. ECaBox workflow and eye maintenance system set-up, including post-mortem eye perfusion and detection in retinal vessels.** (a) Workflow for the resuscitation of the *ex vivo* eye in ECaBox, including rinsing, cannulation and perfusion of porcine and human eye globes, as well as experimental set-up for pressure and flow monitoring, tonometry, retinal fundus imaging, optical coherence tomography, and electrophysiology experiments. Main consumables required, including oxygenated perfusate, corneal, ground and reference electrodes connected to stimulation and detection systems and software are shown. (b) Perfusion pressure and intraocular pressure plots over the first hour of perfusion, showing stabilization after 10 and 20 minutes of perfusion respectively, n=8 pig eyes. (c) Representative OCT B-scan image showing opening of the retinal vessels (pink) and choroidal vessels (yellow) after 5 minutes perfusion, n = 7 pig eyes. (d) Overall retinal thickness is unchanged over the first hour of perfusion, visualised from OCT B-scans, n = 10 pig eyes.

**Fig. S2. Validation of vessel network topology after segmentation.** The centre lines in the reconstructed pig retinal vessels are corrected using a heuristic methodology and displayed for different segment types. Beginning from the raw dataset (left), vessels are segmented and their centre lines identified as described in the text. After this, small cycles (yellow) are detected for further scrutiny, as they are not expected in the scale of vessels resolved by micro-CT. The algorithm identifies spurious segments within the cycles which are marked for removal (red), after which terminal segments are checked for interrupted connections and re-established (purple). Green indicates segments which are retained without modification from the initial extracted network.

**Fig. S3. Perfusion of human donor eyes ex vivo using the ECaBox approach.** Our perfusion strategy was applied to human donor eyes obtained >6 hours post-mortem. (a) Image showing cannulation of ophthalmic artery of human donor eye, representative of (n = 6) human eyes. (b) Placement of the cannula in the ophthalmic artery, (n = 6) human eyes. (c) Retinal blue autofluorescence image (left panel) and fundus imaging (right panel) showing perfusion of white casting material, (n = 3) human eyes. (d) OCT after perfusion with BriteVu casting material. White casting material is visible in the retinal vessels, (n = 4) human eyes.

**Fig. S4. Vessel octant comparisons between early (<5 hours) and late (>10 hours) post-mortem eyes.** (a-b) Schematic of the partition of the globe into octants and bullseye representation. (a) total vessel lengths and (b) vascular volumes expressed as density over octant surface area. (*n* = 3) pig eyes per group. Violin plot colours: light blue = early <5 hours post-mortem, navy = late >10 hours post-mortem. Data are compared by t-test *p≤0.05, **p≤0.01.

**Fig. S5. Retinal pigment epithelium morphology after different post-mortem intervals.** (a) Representative images of phalloidin-stained RPE sheets from eyes stored for 3-, 10- and 24-hours post-mortem at room temperature. (b) Quantification of the RPE cell area based on Phalloidin staining and cell-pose segmentation. Data analysed by one-way ANOVA, (*n* ≥ 5) pig eyes per group, each point represents the average RPE cell area in one eye.

**Fig. S6. Artificial intelligence-based severity grading.** (a) accuracy of EfficientNet-B0 architecture across the 5 folds allows classification of most images. (b) Gradient-weighted class activation mapping (Grad-CAM++) visualises the regions of highest importance for classification decisions of EfficientNet-B0 model.

**Fig. S7. Retinal structural changes in cold versus room temperature conditions.** Representative Haematoxylin and Eosin staining of pig eyes stored for 3-, 10-, 24-hours at (a) 24 ⁰C or (b) 4 ⁰C. (c) Comparison of degeneration severity, with data compared by one-way ANOVA. ns: P > 0.05, *P ≤ 0.05, **P ≤ 0.01, ***P ≤ 0.001, ****P ≤ 0.0001. (*n* ≥ 3) pig eyes per group, each point on the violin plot indicated the average result from the central retina in one eye. Violin plots of non-perfused eyes stored at 4 ⁰C (mint) and at 24 ⁰C (turquoise) are shown. RGC = retinal ganglion cell layer, INL = inner nuclear layer, ONL = outer nuclear layer. Note that the data shown here in (a) is also shown in the Fig. 4 and is included here for comparative purposes with 4 ⁰C-maintained pig eyes.

**Fig. S8. Retinal viability and protein expression in non-perfused post-mortem eyes at 4 ⁰C.** (a-d) Proteomics analyses of porcine retina after different post-mortem intervals at 4 ⁰C. (a) Heatmap showing overall retinal protein changes between 3-,10- and 24-hours post-mortem. The colours in the heatmap represent the abundance of each protein of the proteome quantified in each sample (lower expression shown in blue, higher expression in red). Heatmap representing protein relative abundances (rows) in each replicate and condition (columns). Relative protein abundances have been transformed to z-scores by column, and proteins grouped by hierarchical clustering. (b-d) Data are logarithmic converted and compared with two-way ANOVA with multiple comparisons and also with students t-test. (*n* ≥ 4) pig eyes per group. Analysis of proteomic dataset for (b) photoreceptor markers (c) neuronal markers and (d) cell death markers were examined. ns: p>0.05. (e-g) Single cells disaggregated from retinas, stored in intact eyes at 4 ⁰C for various post-mortem intervals, were examined using the Countess Cell counter. (e) Cell viability count from countess, expressed as a percentage of total cells. (f) Cell size of live cells are compared between 3- and 24-hours post-mortem. (g) Flow cytometry of calcein dye marking live cells, and DAPI marking dead cells, normalised to the total number of events. (e-g) (*n* ≥ 3) eyes from ≥3 pigs. Data are analysed by t-test, ns: P > 0.05, *P ≤ 0.05, **P ≤ 0.01, ***P ≤ 0.001, ****P ≤ 0.0001.

**Fig. S9. The effect of various parameters on light responsiveness in *ex vivo* perfused eyes.** (a) Intensity-dependent light responses in *ex vivo* porcine eyes, *n* = 4. (b) Light responses appear gradually after the onset of perfusion. Plot focusing on the first 60 minutes, showing light responses after 15 minutes, *n =3* pig eyes. (c) The length of ischemia time post-enucleation versus *ex vivo* ERG responsiveness. (d) *In vivo* ERG amplitudes versus duration of *ex vivo* ERG response, linear regression. (e) The effect of *in vivo* IOP on *ex vivo* ERG responsiveness, n.s: p ≥ 0.05. (f) Comparison of baseline light responses *in vivo* between responsive and unresponsive eyes. (c, e, f) Violin plot colours: magenta = ERG responsive, grey = No ERG response, (*n* ≥ 8) pig eyes per group. Data are compared by t-test n.s: p≥0.05.

**Video S1. Retinal fundus imaging of the post-mortem pig eye, showing re-perfusion of all major vessels and their branches.**

**Video S2. Micro-CT imaging of the vascular casted post-mortem pig eye, showing re-perfusion of the entire eye globe and all its vascular beds.**

**Table S1.** Outcome of training for segmentation AI architectures. Evaluation of predictive performance using trained segmentation models against the test set. Attention U-Net outperformed the other models in Intersection Over Unions (IOU) and Dice score, with a minor increase in Hausdorff Distance (HD) compared to the best performer. Since HD is known to be sensitive to outliers, and Dice score combines precision and recall, Attention U-Net is considered the best overall performing network. Bold lettering indicates the best results.

| Metrics | Models |  |  |  |
| --- | --- | --- | --- | --- |
|  | Attention U-Net | Residual U-Net | U-Net | V-Net |
| Hausdorff distance | 1.334667 | **1.229433** | 1.425243 | 1.665169 |
| IOU | **0.849046** | 0.8457 | 0.838065 | 0.849014 |
| F1-score (Dice) | **0.917434** | 0.91536 | 0.910861 | 0.916731 |
| Precision | **0.930055** | 0.927075 | 0.926726 | 0.906582 |
| Recall | 0.910959 | 0.909861 | 0.90205 | **0.934944** |

**Table S2.** Composition of each time point as predicted by the model (3h, 10h, 24h and 24h-perfusion). Degeneration severity (healthy/mild/moderate/severe) is compared, using a Chi-square test. p value > 0.05 indicates a non-significant difference in composition.

|  | Chi-Square test (pairwise comparisons, p-values) |  |  |  |
| --- | --- | --- | --- | --- |
| Time | 3 | 10 | 24 | 24-perf |
| 3 |  | 1.19E-03 | 1.18E-08 | 7.86E-02 |
| 10 | 1.19E-03 |  | 0.043 | 0.27 |
| 24 | 1.18E-08 | 0.043 |  | 3.63E-03 |
| 24-perf | 7.86E-02 | 0.27 | 3.63E-03 |  |

**Table S3.** Electrophysiology responses of *ex vivo* eyes. Ischemia time represents the time elapsed in minutes between enucleation and reperfusion. The *ex vivo* duration recorded here was the time at the last recording, although due to limited availability of the surgical facilities, in some cases eyes could not be followed until they stopped responding. *n* = 15 pig eyes.

| Ischemia time (no perfusion-min) | Ex vivo ERG amplitude (Max-Min μV) | Ex vivo ERG duration (from enucleation) |
| --- | --- | --- |
| 5 | 15.51-5.08 | 76min |
| 15 | 14.79-3.67 | 149min |
| 8 | 15.83-3.67 | 149min |
| 13 | 18.28-7.16 | 120 min |
| 13 | 18.28-7.16 | 78min |
| 15 | 19.23-2.85 | 244min |
| 16 | 20.37-12.38 | 188 min |
| 14 | 26.72-7.8 | 90 min |
| 11 | 16.42-13.43 | 57min |
| 15 | 23.73-11.77 | 45min |
| 14 | 18.82-8.30 | 622min |
| 20 | 36.66-7.62 | 587min |
| 6 | 20.32-2.27 | 801min |
| 9 | 20.96-6.08 | 81min |
| 10 | 25.90-5.12 | 33min |

**Table S4.** Medical profiles and immediate cause of death for human eye donors. Eyes from donors ineligible for cornea donation of advanced age and with comorbidity were obtained within 6-10 hours after death, following informed consent.

| Age | Sex | Cause of medical contraindication for tissue donation | Immediate cause |
| --- | --- | --- | --- |
| 74 | M | Cognitive decline | Asystolic cardiorespiratory arrest |
| 75 | M | Other | Septic shock |
| 90 | M | Age | Ventricular fibrillation |
| 83 | F | Cognitive decline | Respiratory insufficiency |
| 93 | M | Hematological<br>Neoplasia | Septic shock with refractory hypotension and multiple organ failure. |
| 91 | F | Age | Respiratory insufficiency |
| 91 | M | Age | Massive cerebral hematoma (left occipito-parieto-temporal) with mass effect and midline shift. |
| 100 | M | Age | Respiratory insufficiency |
| 89 | F | Cognitive decline | Multi-Organ Failure |
| 94 | M | Age | Acute respiratory insufficiency |
| 97 | M | Age | Asystolic cardiorespiratory arrest |
| 97 | M | Age | Respiratory insufficiency |
| 69 | M | Age | Sepsis |
| 93 | F | Age | Acute respiratory insufficiency |
| 66 | M | Other | Eutanasia |
| 92 | M | Age | Cardiorespiratory arrest |
| 70 | F | COVID | Respiratory insufficiency |

