## Supplementary figures and images for "Retinal resuscitation in post-mortem eyes"

### Figure.S1

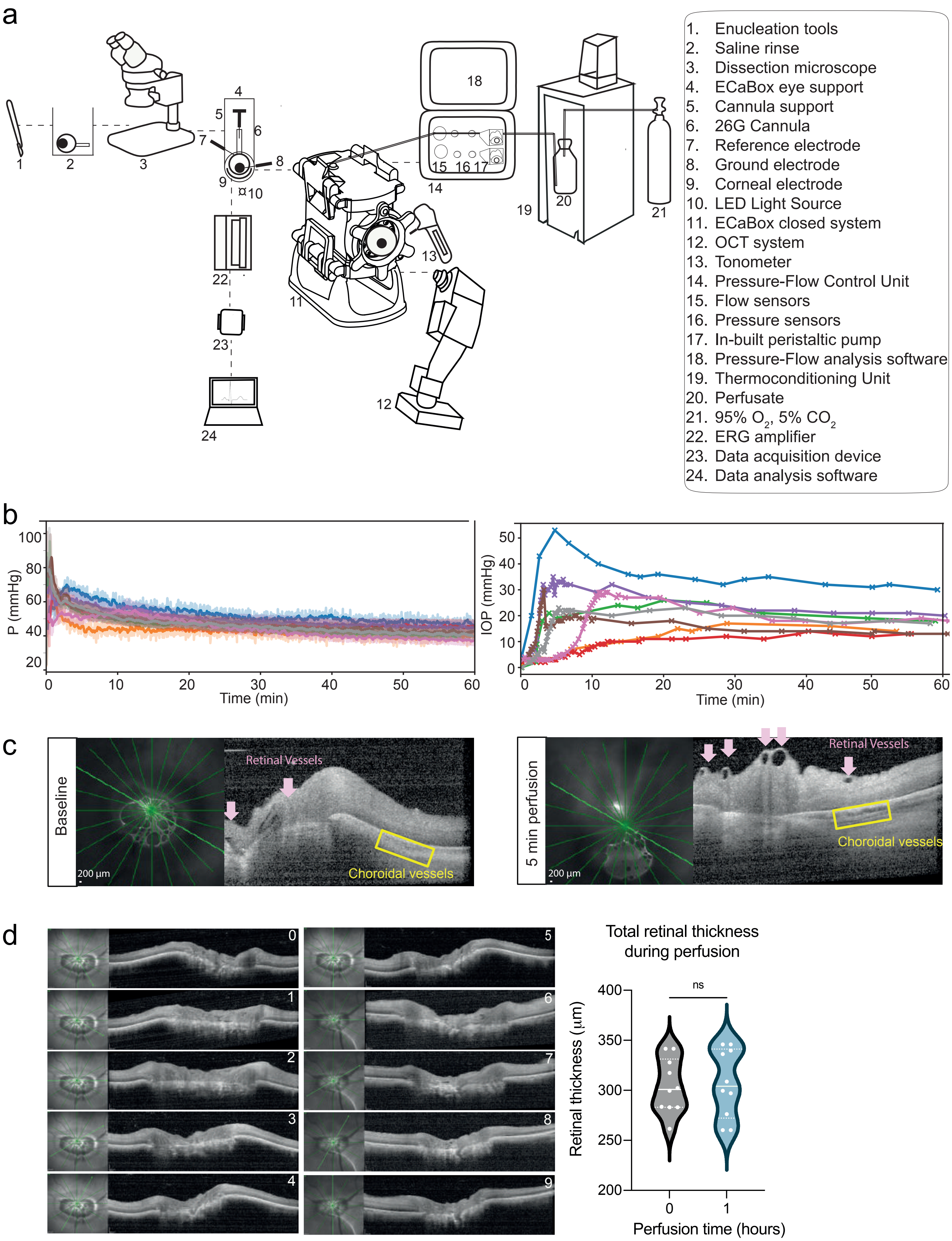

### Figure.S2

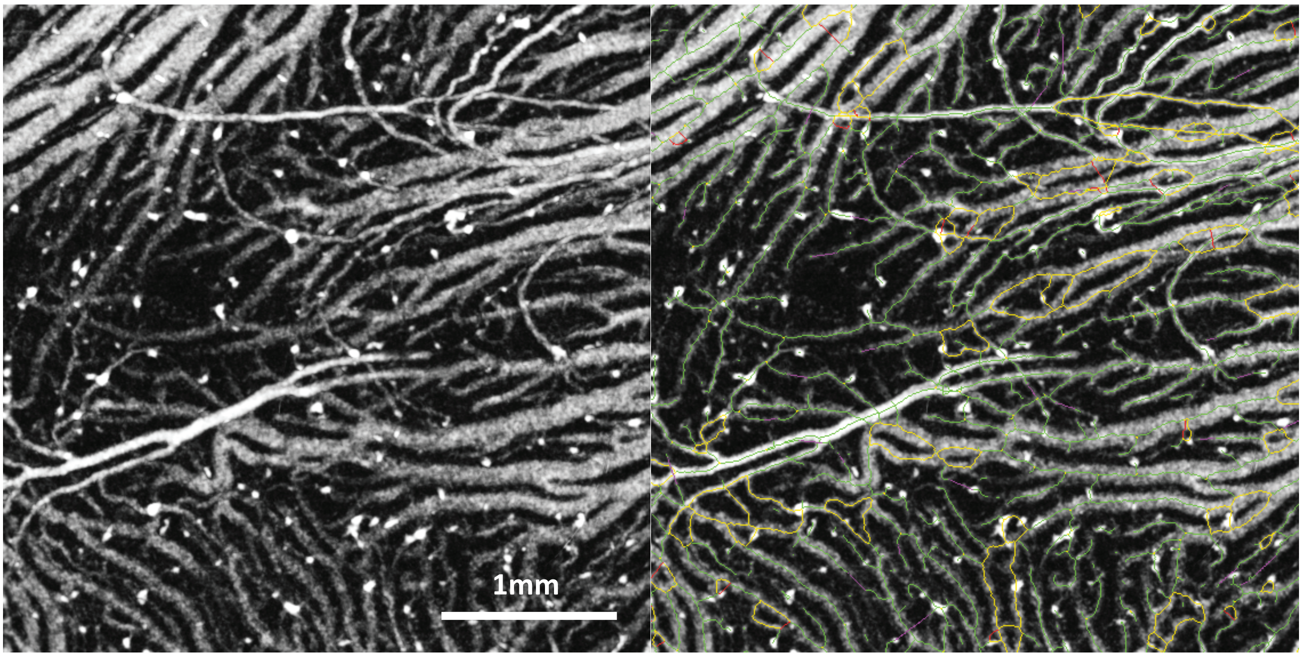


Fig. S2

### Figure.S3

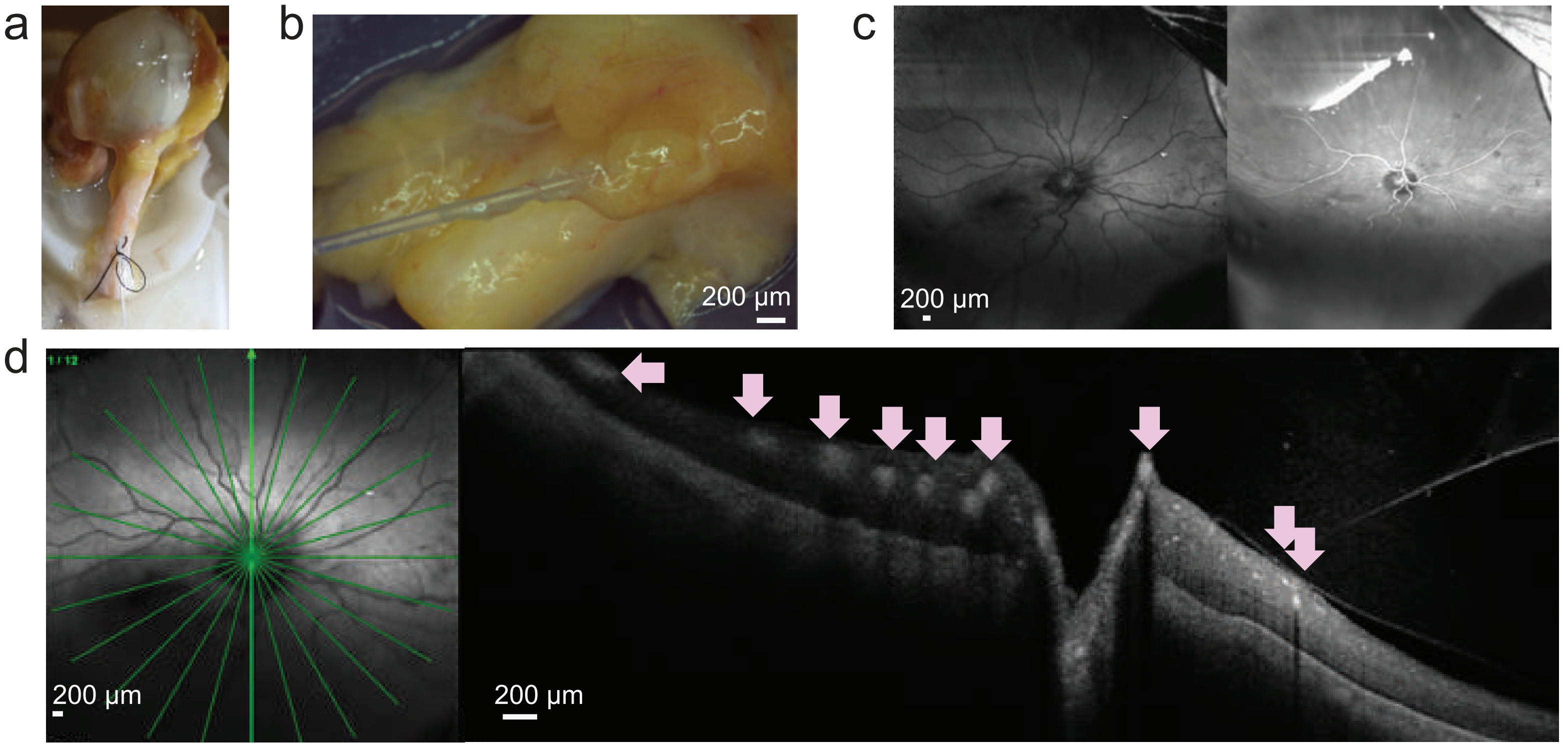


Fig. S3

### Figure.S4

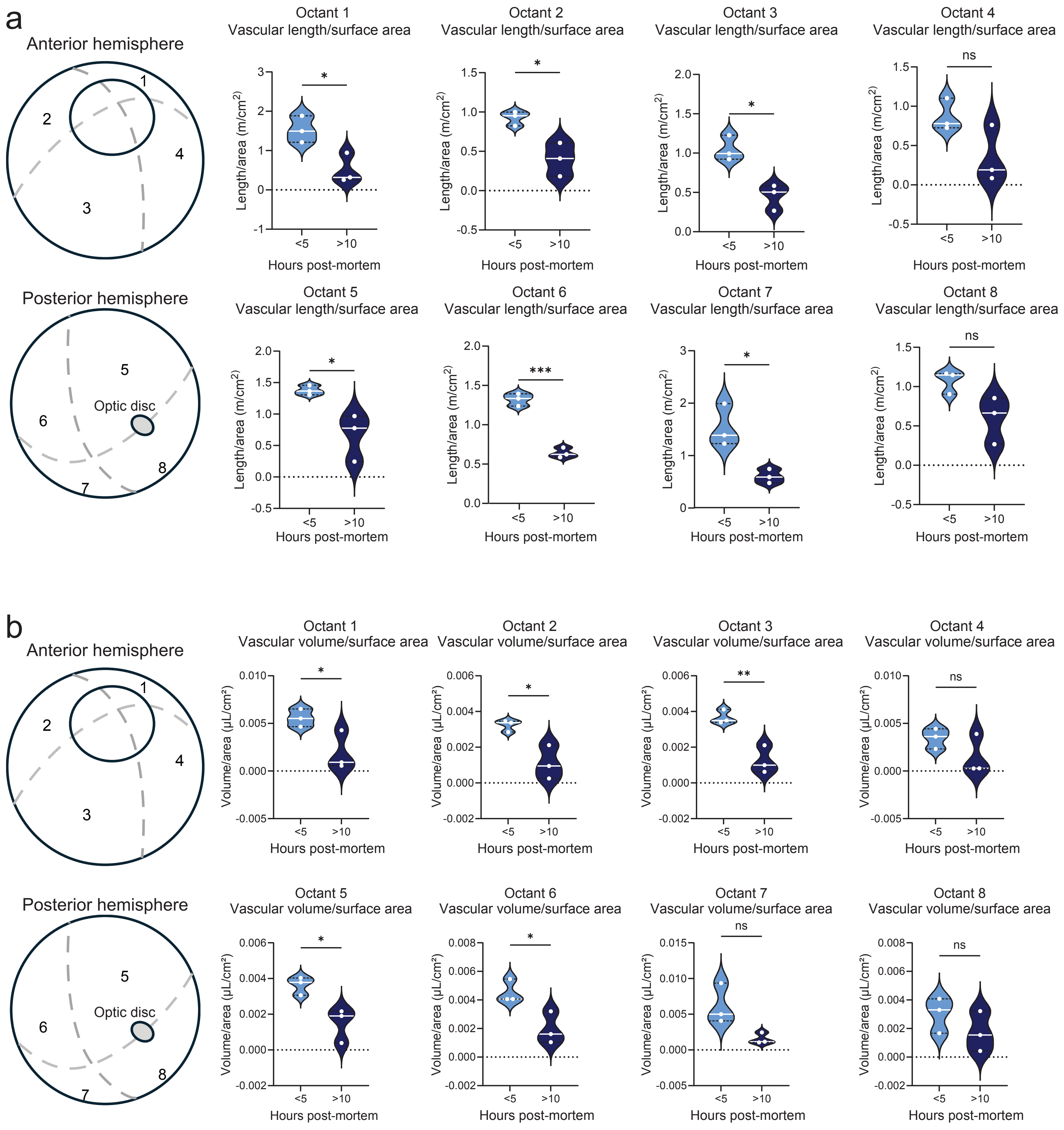

### Figure.S5

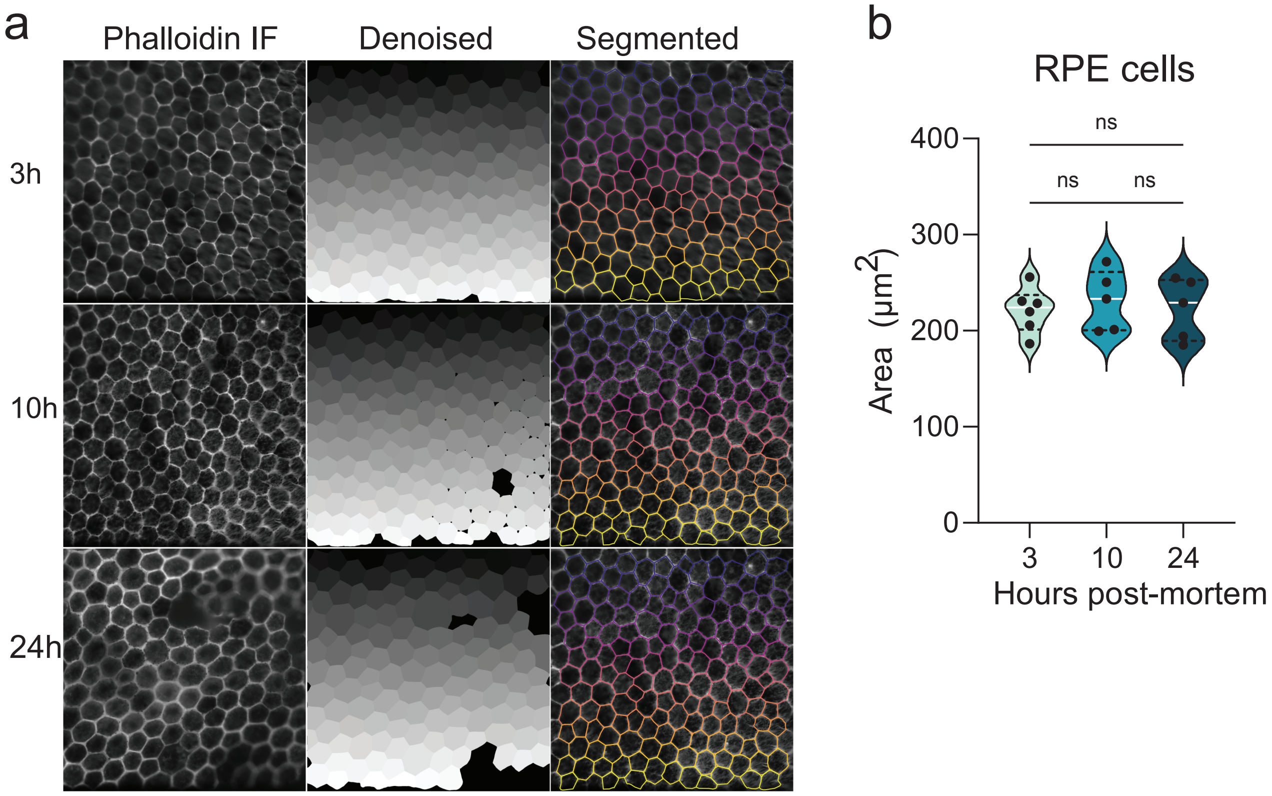

### Figure.S6

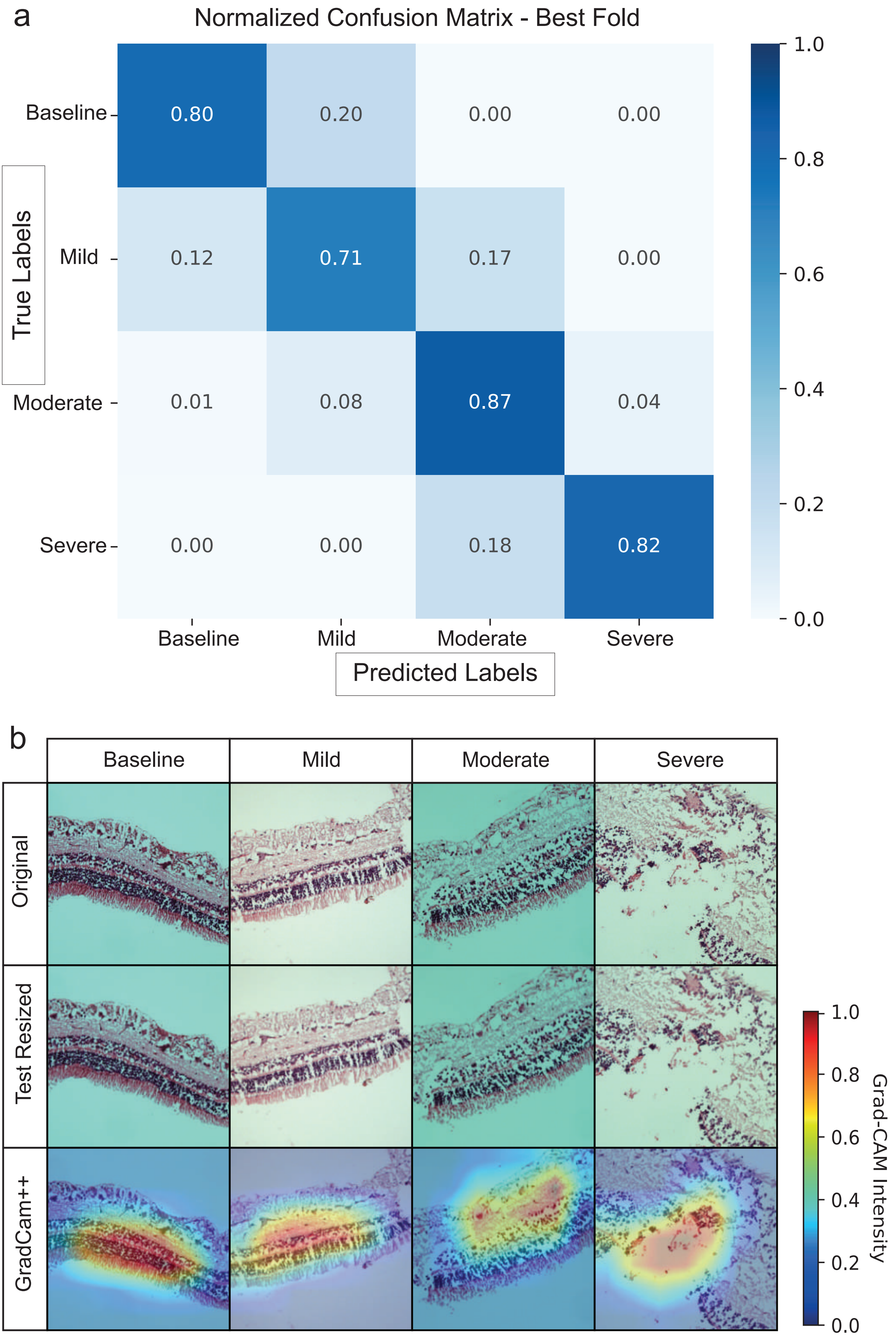

### Figure.S7

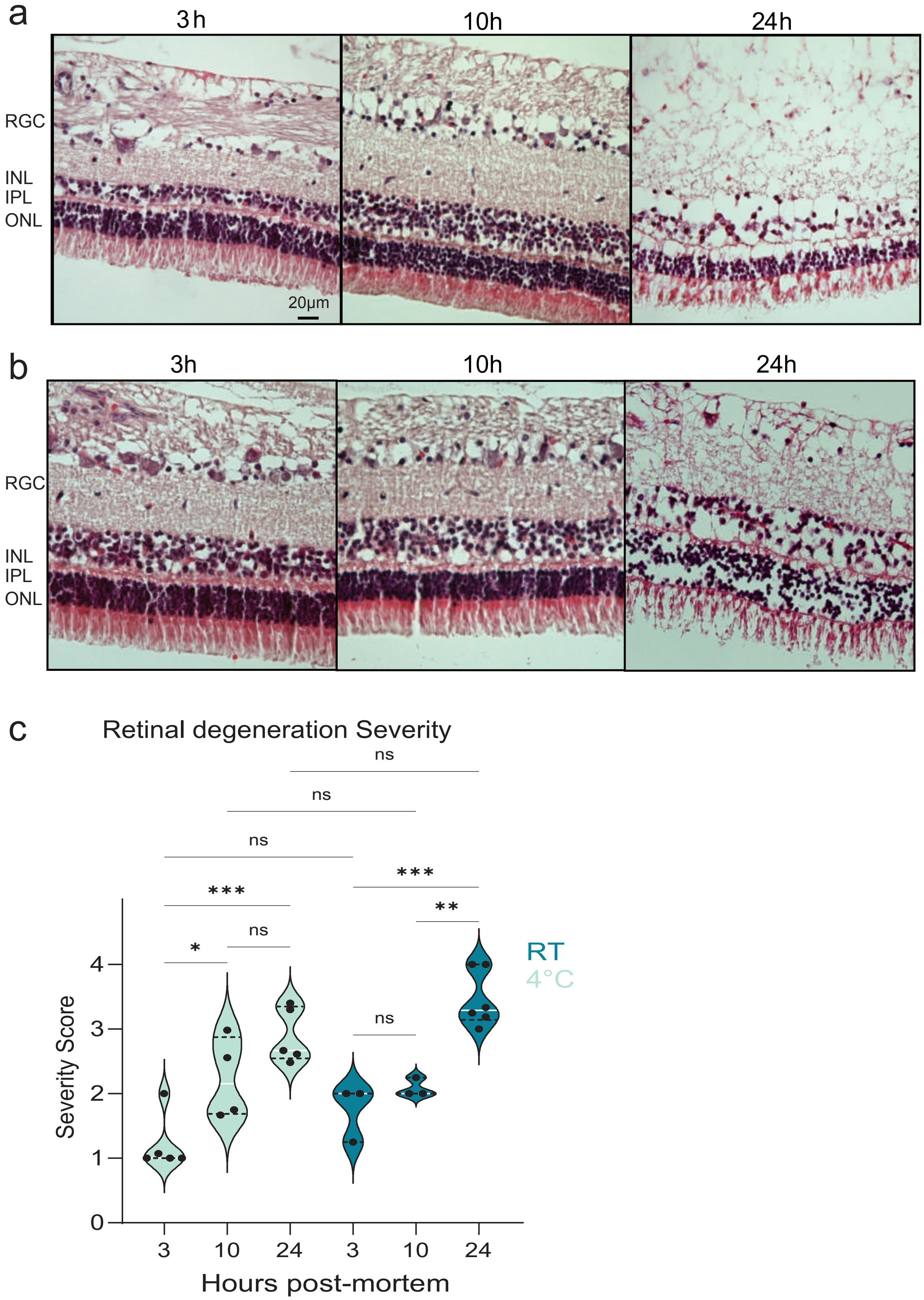

### Figure.S8

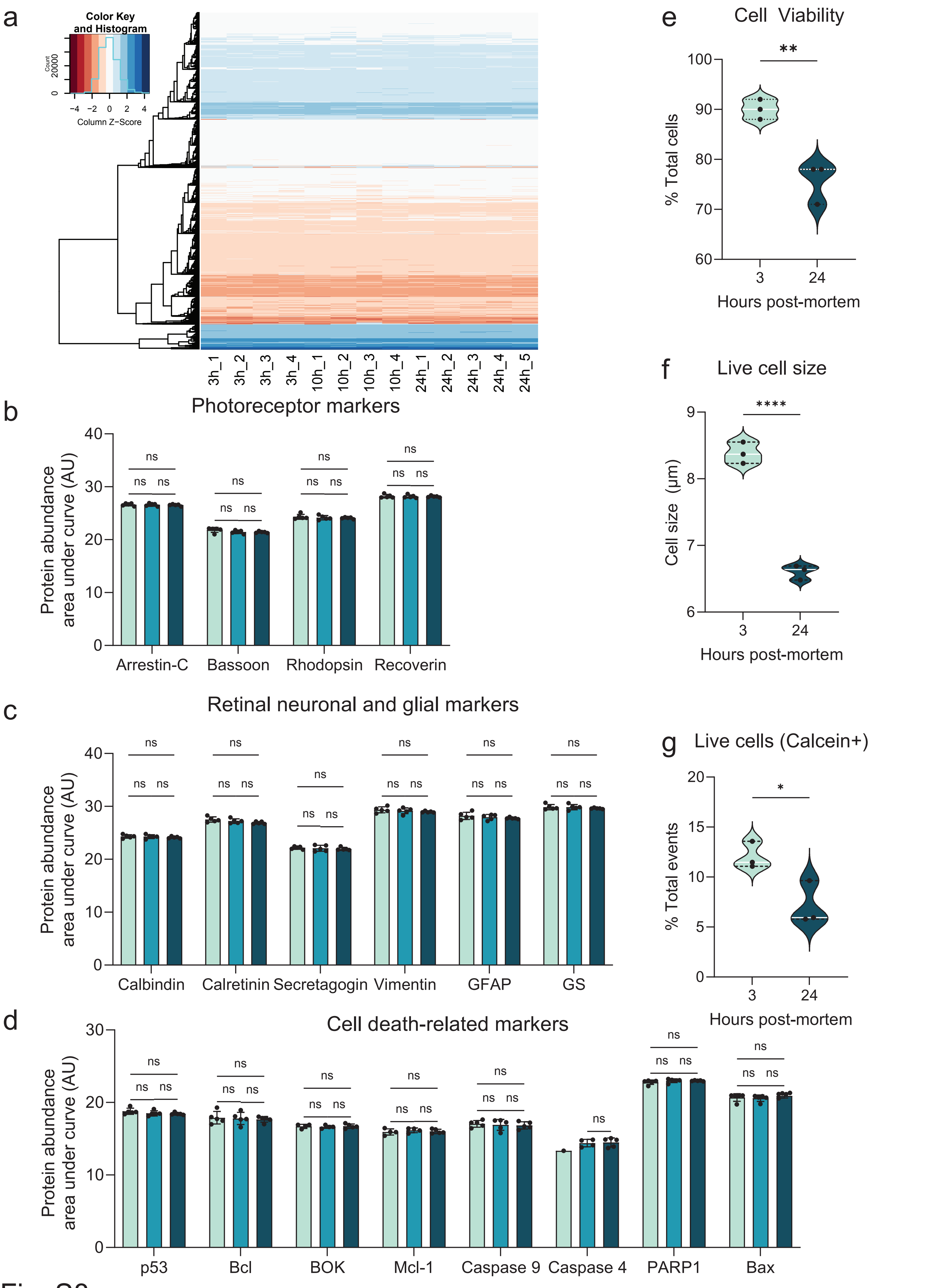

### Figure.S9

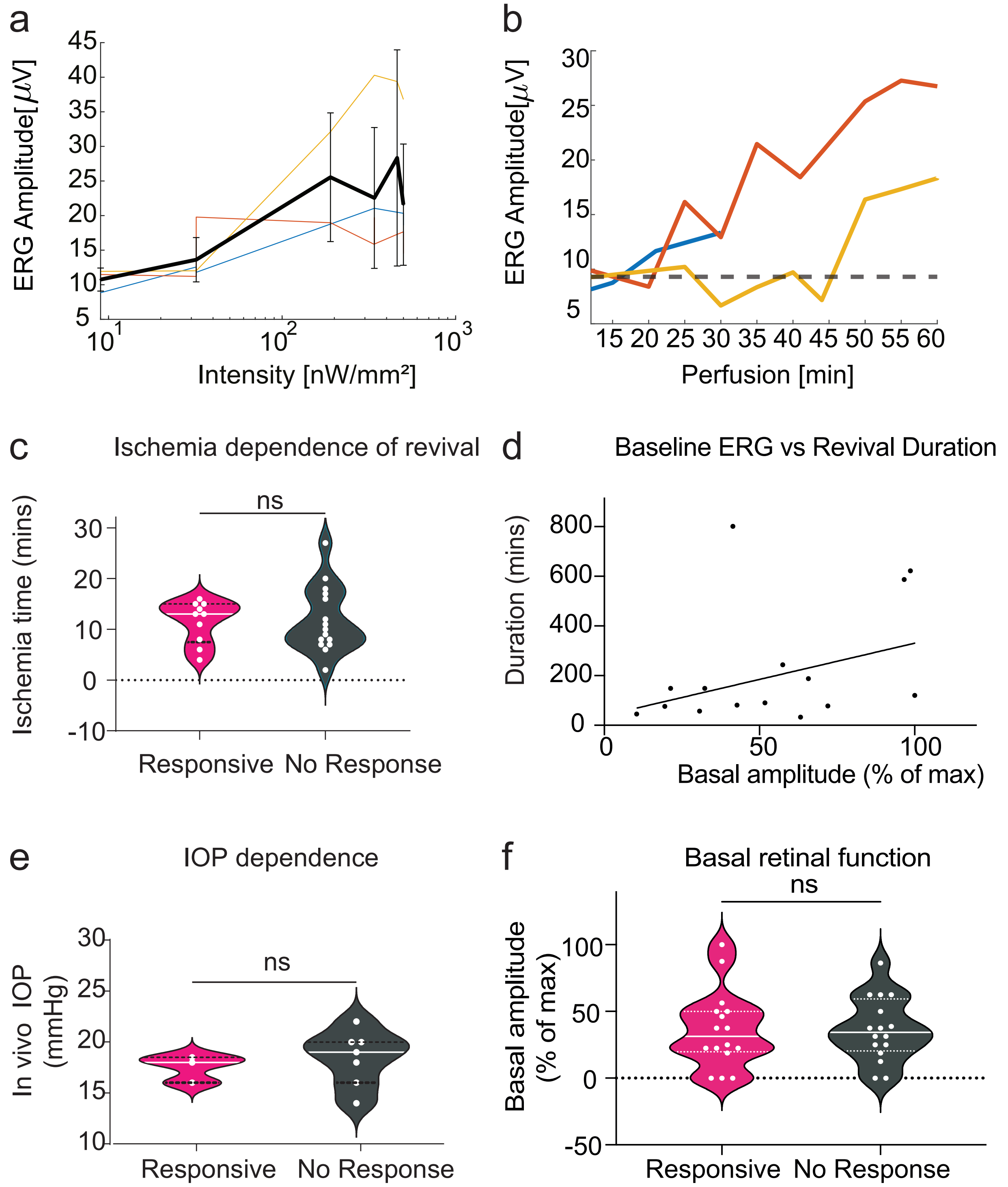
